# SARS-CoV-2 spike D614G variant exhibits highly efficient replication and transmission in hamsters

**DOI:** 10.1101/2020.08.28.271635

**Authors:** Bobo Wing-Yee Mok, Conor J. Cremin, Siu-Ying Lau, Shaofeng Deng, Pin Chen, Anna Jinxia Zhang, Andrew Chak-Yiu Lee, Honglian Liu, Siwen Liu, Timothy Ting-Leung Ng, Hiu-Yin Lao, Eddie Lam-Kwong Lee, Kenneth Siu-Sing Leung, Pui Wang, Kelvin Kai-Wang To, Jasper Fuk-Woo Chan, Kwok-Hung Chan, Kwok-Yung Yuen, Gilman Kit-Hang Siu, Honglin Chen

## Abstract

SARS-CoV-2 causes disease varying in severity from asymptomatic infections to severe respiratory distress and death in humans. The viral factors which determine transmissibility and pathogenicity are not yet clearly characterized. We used the hamster infection model to compare the replication ability and pathogenicity of five SARS-CoV-2 strains isolated from early cases originating in Wuhan, China, in February, and infected individuals returning from Europe and elsewhere in March 2020. The HK-13 and HK-95 isolates showed distinct pathogenicity in hamsters, with higher virus titers and more severe pathological changes in the lungs observed compared to other isolates. HK-95 contains a D614G substitution in the spike protein and demonstrated higher viral gene expression and transmission efficiency in hamsters. Intra-host diversity analysis revealed that further quasi species were generated during hamster infections, indicating that strain-specific adaptive mutants with advantages in replication and transmission will continue to arise and dominate subsequent waves of SARS-CoV-2 dissemination.

## Introduction

A newly emerged β-coronavirus, SARS-CoV-2, which causes COVID-19 disease in humans, attained cross species transmission through a process yet to be defined in detail (Andersen et al., 2020; Wu et al., 2020; Zhou et al., 2020a). Human cases were first identified in Wuhan, China, in December 2019 and SARS-CoV-2 subsequently disseminated worldwide leading to the announcement of a global pandemic by the World Health Organization on March 11, 2020 (Mahase, 2020). More than 20 million laboratory-confirmed cases and over 700,000 deaths have been recorded globally to date (https://coronavirus.jhu.edu/map.html) (Dong et al., 2020). In contrast to SARS-CoV and MERS-CoV, a significant number of SARS-CoV-2 infections are asymptomatic. However, in areas high virus activity a substantial portion of infections lead to severe disease or death (Chen et al., 2020b). While aging and certain underlying medical conditions may predispose individuals to increased severity of COVID-19 disease (Zhang et al., 2020c), it is not clear if viral factors may also contribute to the variable pathogenicity of SARS-CoV-2 in humans (Becerra-Flores and Cardozo, 2020). It is expected that SARS-CoV-2 will continue to be transmitted among humans globally, leading to the emergence of more phenotypic variants. It is therefore important to define the viral factors associated with transmissibility and pathogenicity of SARS-CoV-2. Analysis of a mutant virus with a deletion at the spike protein S1/S2 junction showed that the PRRA polybasic cleavage site is associated with heightened pathogenicity in the model (Lau et al., 2020b). SARS-CoV-2 is of zoonotic origin and is currently in the process of becoming more adapted to humans as it circulates and acquires adaptative mutations. Different variants have already been identified among clinical specimens and in cultured isolates (Gong et al., 2020; Su et al., 2020). Some deletion or mutation variants may not be recognized by general sequencing protocols, for which the readout only shows the dominant population in clinical specimens or cell culture samples, but can be detected by more sensitive methods (Wong et al., 2020).

SARS-CoV-2 was first identified in China in December 2019 (Wu et al., 2020; Zhou et al., 2020a). A response adopting aggressive control measures, including the lock down of a city of 10 million people (Wuhan) and the wider Hubei Province (population: 56 million) in January 2020, significantly limited the further dissemination of the virus within China (Ji et al., 2020; Lau et al., 2020a). SARS-CoV-2 has been efficiently transmitting among humans since it was first recognized, according to studies from the early outbreak (Chan et al., 2020a; Liu et al., 2020). Subsequent transmission of SARS-CoV-2 in Europe, the US and other countries has resulted in more widespread human infections since March 2020. As more humans are exposed to the SARS-CoV-2 virus, it is expected that more host adapted phenotypic variants of the virus will emerge. It is important to determine whether some emerging variants may have the potential to go on to become the dominant strain in the coming waves of circulation. Indeed, a strain bearing a D614G mutation in the spike protein, first observed in January 2020 among isolates from China and Europe, has since become the dominant population in the recent transmissions occurring in Europe and the US (Korber et al., 2020). Studies using pseudo-viruses containing spike genes derived from natural isolates have shown that variants harboring D614G infect cells more effectively (Daniloski et al., 2020; Zhang et al., 2020b). Although D614G is in the spike protein, the D614G variant is still susceptible to neutralization by antibodies raised against strains lacking this mutation (Korber et al., 2020). To further understand the properties of the D614G variant, we analyzed pathogenicity, virus replication efficiency and the global transcriptome of the host response in the airways of SARS-CoV-2 D614G variant infected hamsters. Although it is not clear if the D614G variant causes more severe disease in humans, our data showed that SARS-CoV-2 containing D614G replicates more efficiently and causes more severe pathological changes in the lung tissues of infected animals when compared to isolates lacking this mutation.

## Results

### SARS-CoV-2 genomic variants exhibit variable pathogenicity in hamsters

Five isolates were selected to study variability in the pathogenicity of SARS-CoV-2 strains and the host response to such variants in a hamster infection model. These strains represent isolates from the early outbreak in China (HK-8, HK-13 and HK-15) and subsequent outbreaks in Europe and elsewhere (HK-92 and HK-95) (Table S1). Phylogenetic analysis revealed that these strains belonged to distinctive GISAID phylogenetic clades (**Figure S1**). The sequences of these five isolates were compared with that of the index isolate (Wuhan-Hu-1), characterized during the early outbreak in Wuhan city, China (Wu et al., 2020; Zhou et al., 2020a), revealing a range of variations in the untranslated region and *Orf1a, Orf1b, Orf3, Orf8, N* and *S* genes among strains (**Table 1**). HK-95 is an isolate characterized from a traveler returning to Hong Kong from Egypt and carries a D614G substitution in the spike gene (**Figure S1**). Hamsters are a highly susceptible model for studying SARS-CoV-2 infection, with disease in hamsters closely simulating COVID-19 disease in humans (Chan et al., 2020b). Infection progresses rapidly in hamsters, but infected animals then recover after about one-week of infection. We infected hamsters with these five strains of SARS-CoV-2 and monitored disease presentation for 5 days. All five isolates caused significant body weight loss compared to the control group (**Figure 1A**). In SARS-CoV-2-infected hamsters, three major histopathological changes were observed in lungs: various degrees of bronchial or bronchiolar inflammation (bronchiolitis), lung parenchymal inflammatory damage (alveolitis) and pulmonary blood vessel inflammation (vasculitis). These pathological changes were most severe at day 5 after virus inoculation. The HK-13 and HK-95 isolates caused more extensive bronchiolar cell death, diffuse alveolar space exudation and infiltration and lung consolidation, compared to HK-8 (**Figure 1B**). In our previous report, at day 4 post-infection (pi) with HK-001a virus, alveoli had already started to show focal areas of cell proliferation indicating resolution of inflammation (Chan et al., 2020b; Lau et al., 2020b). However, such indications of resolution were not seen in HK-95- or HK-13-infected hamsters, even at day 5 post-infection, indicating that the acute lung inflammation caused by both of these strains may last longer than that triggered by infection with other SARS-CoV-2 strains.

**Figure 1.**
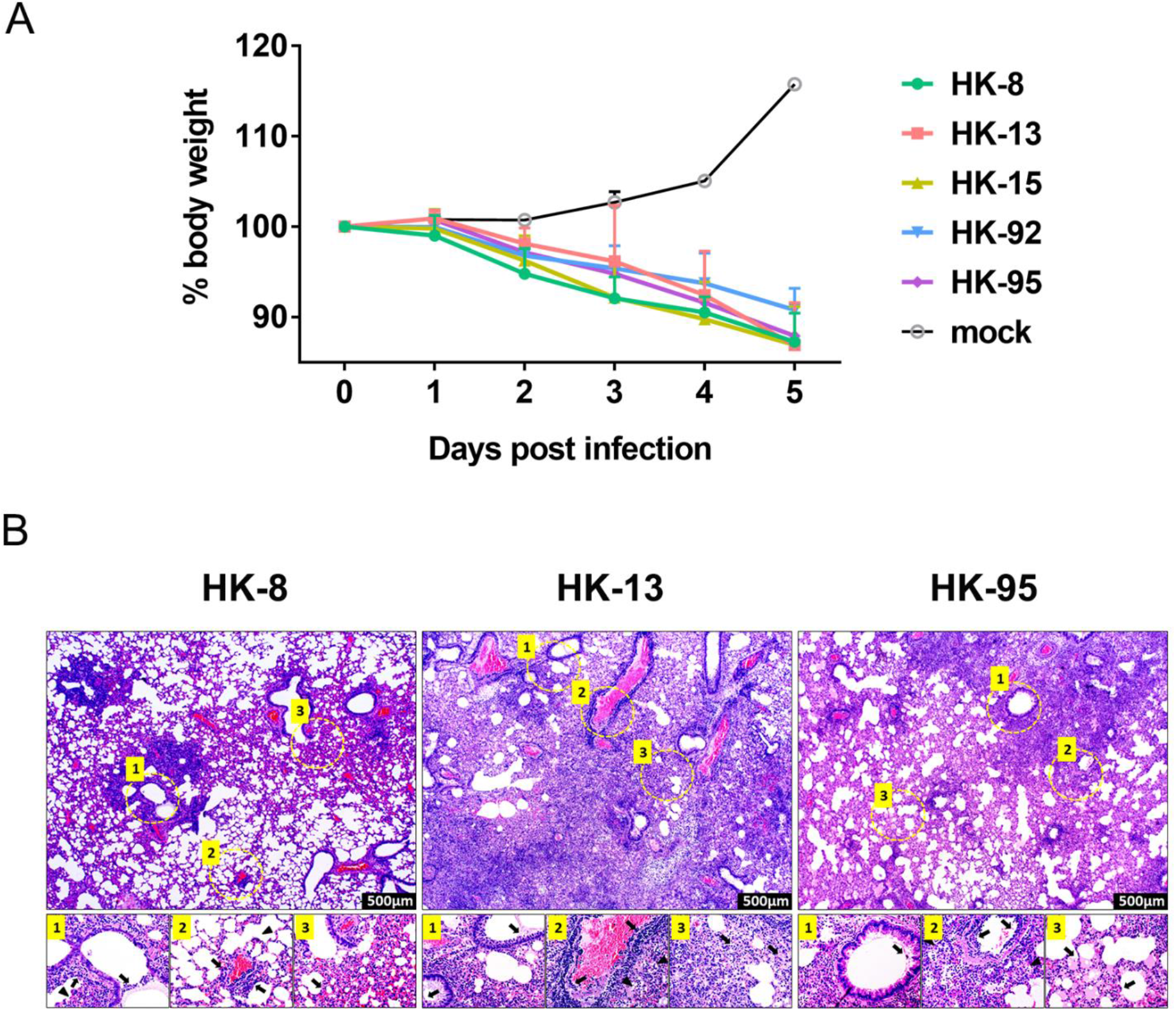
Body weight and histopathological changes in golden Syrian hamsters challenged with different strains of SARS-CoV-2. (A) Body weight change in hamsters after viral infection. Body weights of virus-infected and mock-infected hamsters (n=3) were monitored daily for 5 days. Data are shown as the mean ± SD percentages of the starting weight. (B) Haemotoxylin and eosin (H&E) staining of lung sections from SARS-CoV-2 (HK-8, HK-13 and HK-95) infected hamsters, collected at 5 days post-infection (dpi). The lower photographs depict higher magnification images of the regions denoted by circles in upper photographs (upper photographs = 500x magnification). **HK-8**: Upper panel shows patchy inflammatory damage with areas showing peribronchiolar and perivascular infiltration, blood vessel congestion and areas of alveolar wall thickening, while alveolar exudation and infiltration were not apparent. Lower panel: (1) bronchiolar luminal exudation with cell debris (arrows), surrounding alveolar wall infiltration (arrowhead); (2) a small sized blood vessel shows perivascular and endothelial infiltration (arrows), but the surrounding alveolar space shows no infiltration or exudation (arrowhead); (3) alveolar wall capillary congestion and moderate infiltration (arrow). **HK-13**: Upper panel shows diffuse lung tissue inflammatory damage with massive alveolar space infiltration and exudation; perivascular infiltration and endothelial infiltration can be observed in all blood vessels in this lung section. Lower panel: (1) mild bronchiolar luminal exudation with cell debris (arrows); (2) a large blood vessel shows severe endothelial infiltration and vessel wall infiltration (arrows) and surrounding alveolar space infiltration and exudation (arrowheads); (3) alveolar structure destroyed by septal edema and alveolar space infiltration and fluid exudation (arrows). **HK-95**: Upper panel shows diffuse lung tissue damage with massive alveolar space exudation and infiltration; all blood vessels in the field show perivascular infiltration and endothelial infiltration. A moderate degree of bronchiolar epithelial cell death and luminal exudation is seen. Lower panel: (1) bronchiolar luminal exudation with cell debris (arrow); (2) a medium sized blood vessel shows severe endothelial infiltration and vessel wall infiltration (arrows), with alveolar space infiltration in alveoli surrounding the vessel (arrowheads); (3) alveolar structure has reduced cellularity but shows alveolar septal edema and alveolar space fluid exudation (arrows).

**Table 1.**
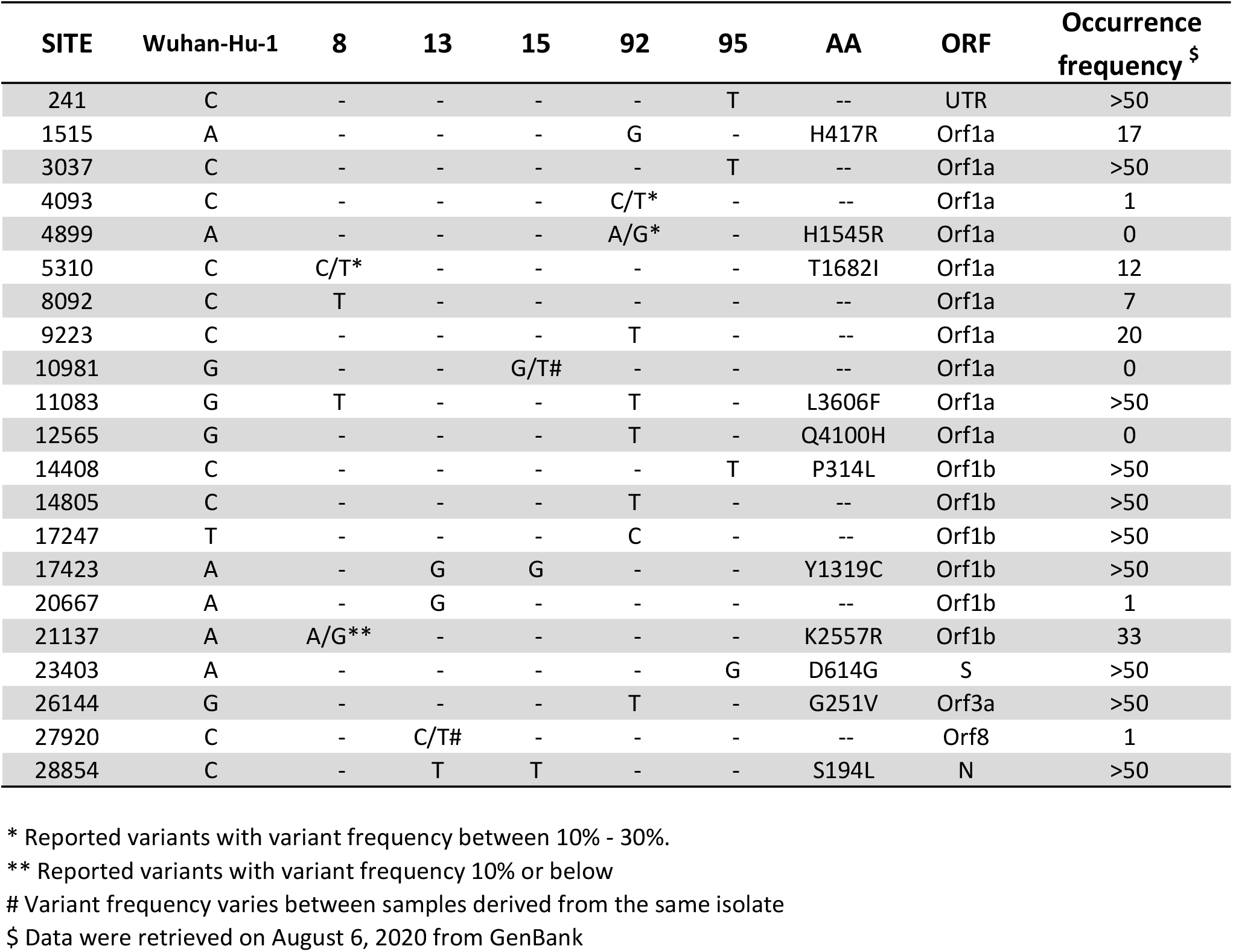
Sequence diversity in SARS-CoV-2 isolates

### Differences in replication rates of SARS-CoV-2 isolates in infected hamsters

Examination of virus titers in lung tissues and nasal washes at day 5 post-infection showed that HK-13 and HK-95 replicate to significantly higher levels than the other three isolates (**Figure 2A and S2**). We then analyzed viral gene expression in the lung tissues of hamsters infected with different isolates using RNA-seq analysis. Consistent with the virus titers, viral gene expression also showed a similar distribution between isolates when comparing the average expression of all viral genes per isolate (black line) to viral titer profiles (**Figure 2B**). Interestingly, the expression distribution between each of the viral genes for each isolate is very similar across all isolates, with differences in overall expression being due to a proportional change in expression across all genes. This indicates that the observed differences in viral gene expression can be attributed to isolates maintaining different rates of replication. These differences in replication suggest that some viral isolates may be more constrained by host-specific factors and as such are unable to achieve an optimal rate of replication, as is seen for HK-8. This also implies that individual isolates interact differently with their hosts during the infection process. To assess if constraints to viral replication are reflected in the severity of the host response induced by SARS-CoV-2 infection, we performed an initial characterization of the hamster host response to infection.

**Figure 2.**
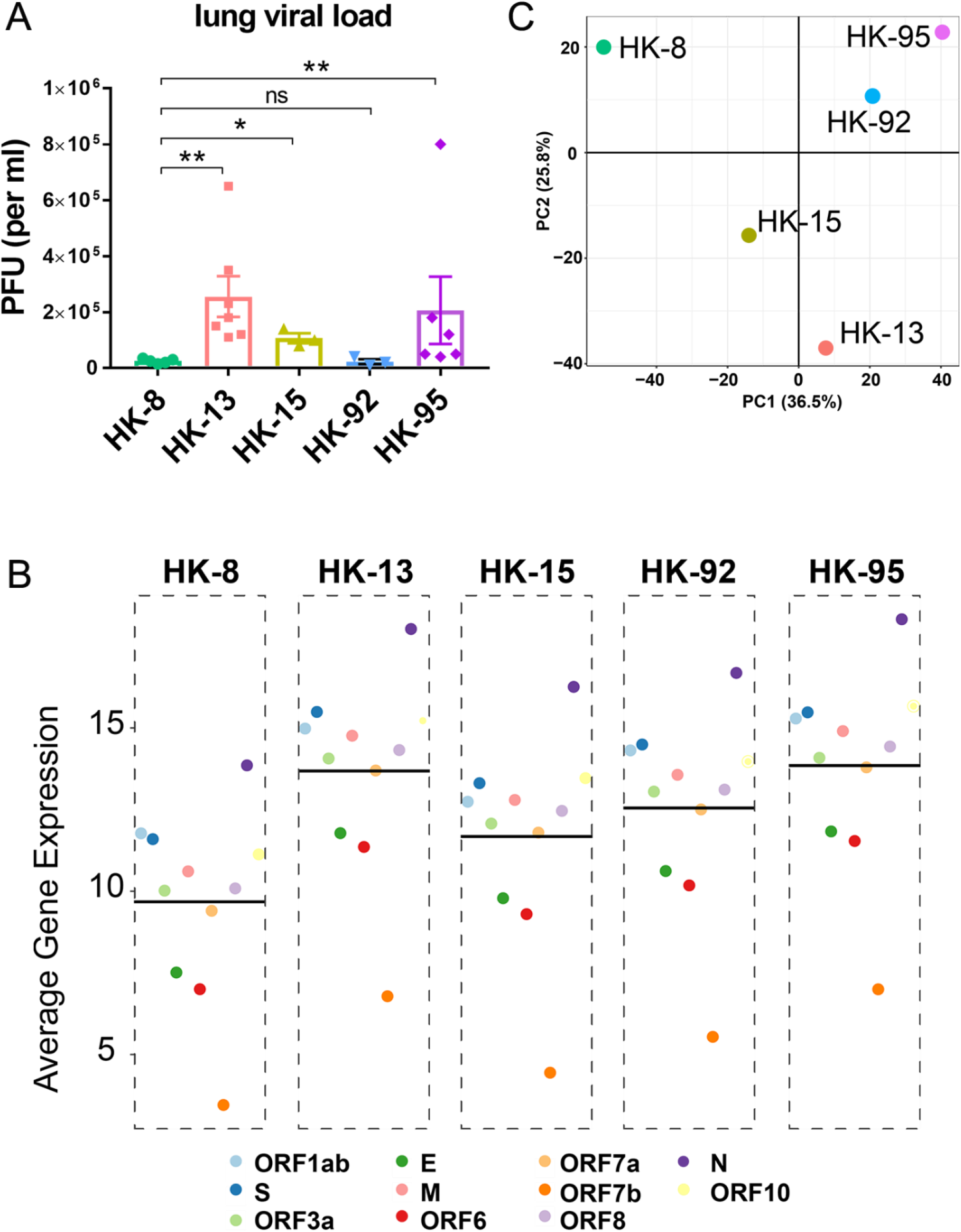
Viral replication and transcriptome response to different SARS-CoV-2 isolates in hamster lungs. (A) Virus replication in hamsters infected with different SARS-CoV-2 isolates. Three to seven hamsters per group were euthanized on day 5 post-infection for viral titration. Virus titers in lungs were determined by plaque assays (PFU/ml) and displayed as mean (± SEM). (B) Average viral gene expression in samples from hamsters infected with each isolate. Horizontal black lines indicate the overall mean of average expression values for viral genes per isolate. (C) Transcriptome response to different SARS-CoV-2 isolates in hamster lungs. Principal component analysis of global transcription profiles of significantly expressed host genes for infections with each isolate, compared to uninfected conditions. Individual data points and means ± SEM are also shown. Statistical significance was calculated by Mann Whitney Wilcoxon Test; * denotes*p<0.05*, ** denotes*p<0.005* and ns denotes *non-significant*.

### Differential expression of SARS-CoV-2 genes and host response in hamsters

To further understand the global host response elicited by difference isolates, we grouped the samples in a principal component analysis (PCA) (Figure 2C). In this PCA space, we observed transcriptional perturbations along the two principal components, both account for more than 25% of sample variation (Figure 2C). This analysis suggested unique expression signatures in hamster host responses to infection with different SARS-CoV-2 isolates. In addition, volcano plots show that viral isolates HK-13 and HK-95, which demonstrate higher rates of replication than isolate HK-8, incited significant upregulation of host gene expression compared to HK-8 (**Figure 3**). Unsurprisingly, the host responses provoked by HK-8 and HK-15 were similar, as these isolates share a similar replication rate. These profiles suggest that higher rates of SARS-CoV-2 replication are a significant driving factor in triggering host responses, which may associate with greater disease severity in humans (Blanco-Melo et al., 2020; Zhou et al., 2020b). However, the differential expression profile generated by comparing HK-8 to HK-92 is similar to that of HK-8 vs HK-95. Since HK-92 has a considerably lower replication efficiency than HK-95 in hamsters (**Figure 2**), this suggests that factors independent of the replication of HK-92 are also involved in inducing the significant upregulation of host gene expression seen here. This offers an explanation as to the close coordinate positioning of HK-92 to HK-95 in our PCA analysis (**Figure 2C**), and why HK-13 is positioned further away, despite possessing a similar replication rate to HK-95 (**Figure 2A and S2**). Our analysis has made it clear that the upregulation of the host response is heavily dependent on the specific viral SARS-CoV-2 isolate that is causing infection. Although viral replication is a significant driver in stimulating the host response in hamsters, SARS-CoV-2 appears to have developed replication-independent strategies to influence responses in infected hosts.

**Figure 3.**
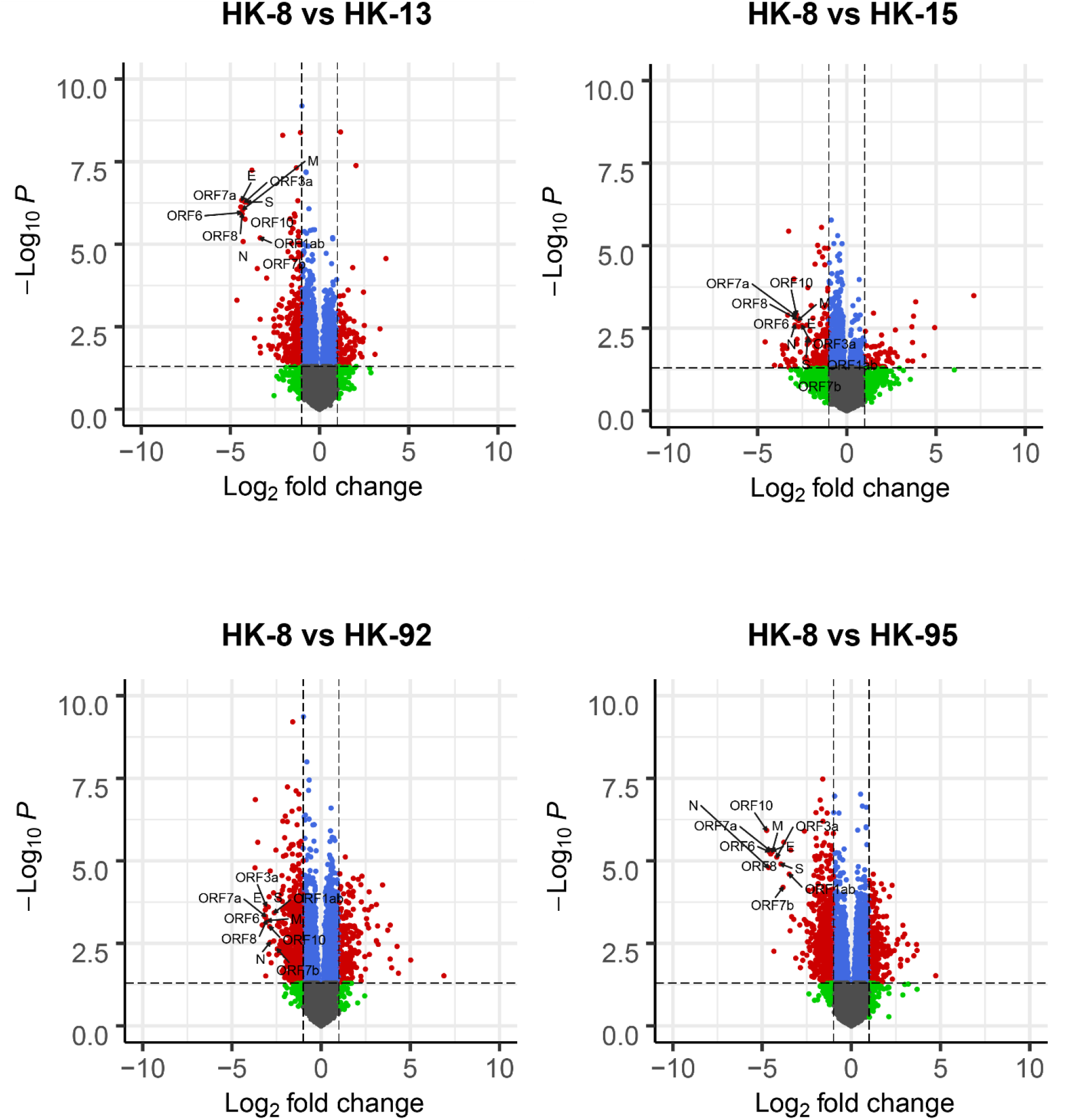
Volcano plots comparing differentially expressed genes (DEGs) in HK-8-infected hamster lungs to those in hamsters infected with HK-13, HK-15, HK-92 or HK-95. DEGs: log2FoldChange ⩾ 1 and p-adjusted value ⩽ 0.05.

Hamsters have proven to be a useful live model for studying the dynamics of viral infection and are now routinely used to assess SARS-CoV-2 pathogenesis (Chan et al., 2020b; Lau et al., 2020b; Zhang et al., 2020a). However, the current annotation that is available to describe gene functionality and perform network analysis in hamsters is limited. Therefore, we opted to use the gene functional annotation of a well-defined animal model within close evolutionary distance of the hamster. Our evolutionary analysis identified mice as being suitably closely related to hamsters (**Figure S3**). Therefore, our gene enrichment analysis of differentially expressed hamster-mouse gene orthologues using mouse Gene Ontology (GO) annotation is anticipated to give a fair representation of the induced host response to SARS-CoV-2, without enrichment of redundant hamster-specific processes. GO enrichment analysis identified many functional groups corresponding to T-cell activation and chromatin remodeling as being the networks most significantly upregulated by SARS-CoV-2 infection, regardless of viral isolate used, when compared to samples from uninfected hamsters (**Figure 4**). Activation of T-cells is indicative of upregulation of the adaptive immune response and the host’s attempts to restrict viral pathogenesis. The upregulation of chromatin remodeling pathways indicates significant disruption to DNA architecture and an increase in nuclear repair processes, potentially due to cellular stress caused by the SARS-CoV-2 virus during infection. We also conducted GO enrichment analysis of differentially expressed genes in hamster lungs induced by the various SARS-CoV-2 isolates, this time conducting comparisons to samples from HK-95 hamster infections. Our result indicates that subverted gene networks are subject to strain specific targeting processes by the more pathogenic strains of SARS-CoV-2, but not by HK-8 (**Figure S4**). Overall, the main effects of SARS-CoV-2 infection involved activation of the T cell response and disruption of regulatory processes in the nuclear microenvironment of infected cells.

**Figure 4.**
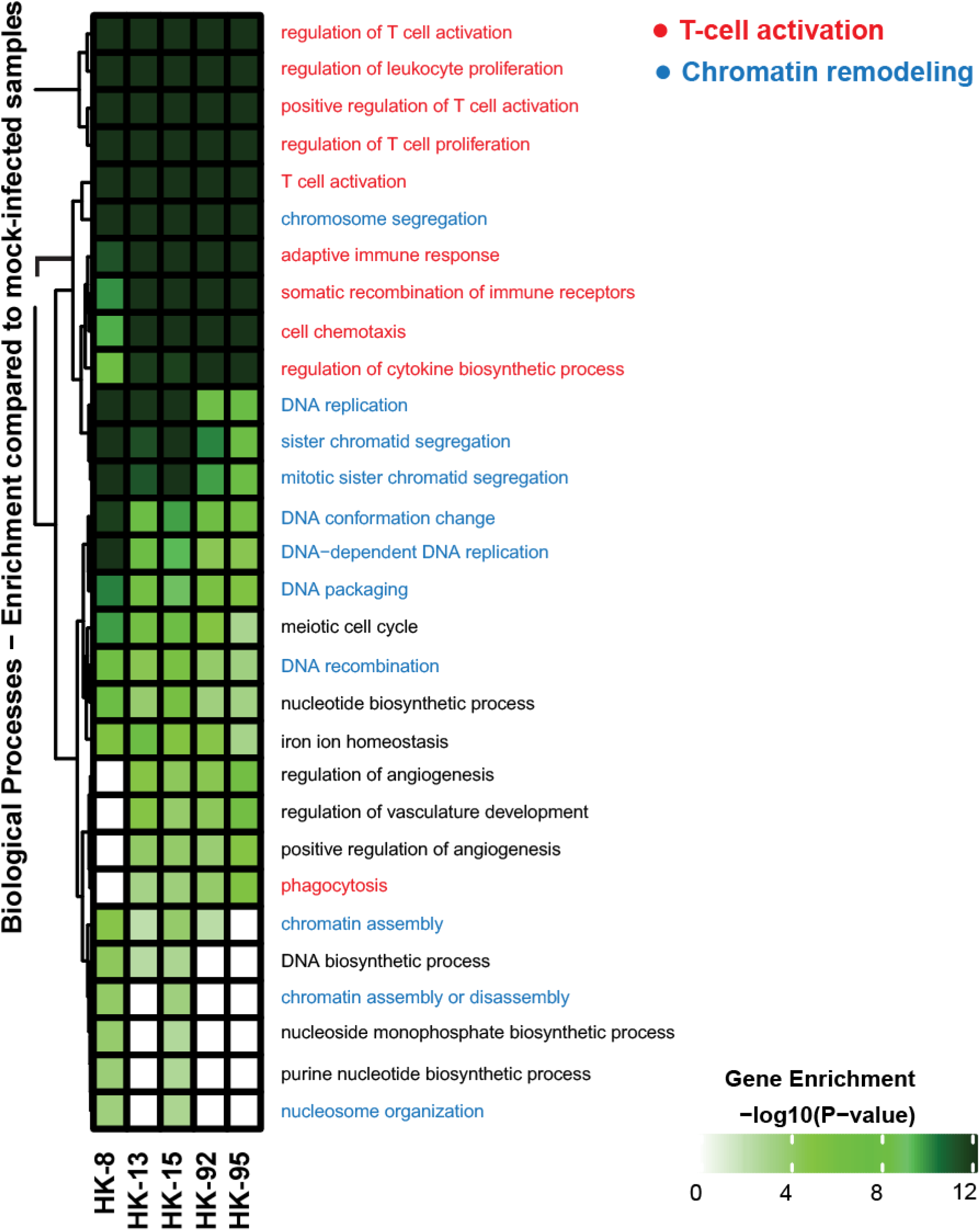
Differential gene expression in SARS-CoV-2 infection. Heatmap depicting the expression levels of enriched gene sets in lungs collected from hamsters infected with the indicated viruses at 5 dpi. Gene enrichment was performed on differentially expressed genes identified in comparisons between infections with each of the 5 different SARS-CoV-2 isolates and uninfected controls. DEGs that were identified as hamster–mouse orthologues were extracted and matched to gene members of mouse GO biological processes. The top 30 GO biological processes with significant enrichment (FDR ≤ 0.05) across infected conditions are displayed.

### D614G variant demonstrates higher transmissibility in hamsters

Surveillance of the evolution of SARS-CoV-2 during circulation in humans has identified various mutations which may relate to the infectivity of virus (Chen et al., 2020a; Korber et al., 2020) The spike D614G variant of SARS-CoV-2 has been the clearly dominant population since March 2020, suggesting that it has enhanced infectivity in humans (Becerra-Flores and Cardozo, 2020; Daniloski et al., 2020; Zhang et al., 2020b). We conducted an experiment using the hamster model to examine if the transmission ability of the HK-95 strain, which carries spike D614G, may be enhanced compared to two other isolates (HK-8 and HK-13) which lack the D614G substitution (**Figure S5**). Notably, higher virus titers were found in the lung and nasal turbinate tissues of recipient hamsters co-housed with HK-95-infected hamsters than in those secondarily exposed to HK-8 or HK-13 (**Figure 5A**), which differs from the direct infection experiment, where HK-13 and HK-95 demonstrated similar viral titers in respiratory tissues **(Figure 2A and S2)**. We did not examine if naive hamsters in the HK-95 group were infected at an earlier time point than those in other groups due to limited availability of experimental animals. But given the higher viral titers in the lung and nasal turbinate tissues, it is possible that hamsters co-housed with a HK-95-infected hamster were infected earlier than those co-housed with hamsters infected with HK-8 or HK-13, and that this facilitated greater virus replication in recipients. It is unclear how the D614G mutation affects virus replication. The efficiency of spike protein cleavage is known to associate with coronavirus replication (Hoffmann et al., 2020; Jaimes et al., 2020). We found that introduction of the D614G mutation into the Wuhan-Hu-1 prototype sequence enhanced SARS-CoV-2 spike cleavage in cells (**Figure 5B**). Since D614G is not located at either of the cleavage sites in the spike protein of SARS-CoV-2, the molecular basis of this effect remains to be investigated. Taken together, these results confirm that acquisition of D614G promotes SARS-CoV-2 infectivity in a mammalian model of SARS-CoV-2 infection.

**Figure 5.**
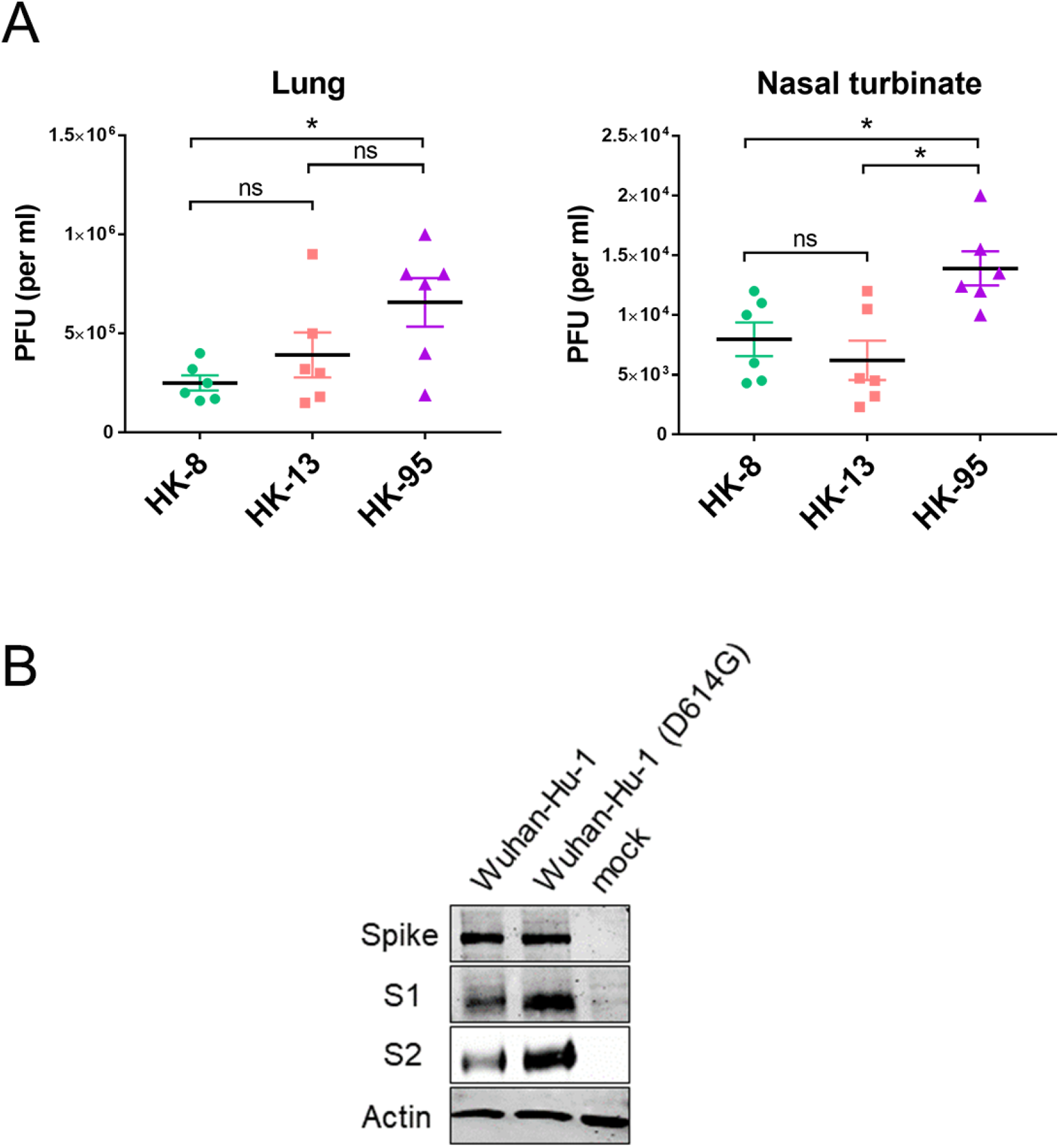
D614G enhances efficiency of spike protein cleavage and transmission of SARS-CoV-2 isolates in hamsters. (A) Transmission of SARS-CoV-2 isolates. For each isolate, two groups of three naïve hamsters were each co-housed with one inoculated donor on day 1 post-infection. Viral loads (PFU/ml) in lungs and nasal turbinates of naïve contact hamsters (n=6 per isolate) at 3 days after exposure are shown. Data are presented as individual data points and means ± SEM. Statistical significance was calculated by Mann Whitney Wilcoxon Test; (*) p value < 0.05, ns – not significant. (B) The human codon-optimized spike gene of Wuhan-Hu-1 SARS-CoV-2 was cloned into the Flag-tagged-pCAGEN vector. A pCAGEN-S-D614G-Flag mutant was constructed using a QuikChange site-directed mutagenesis kit (Agilent) according to the standard protocol. Protein expression and cleavage in transfected HEK293T-ACE2 cells was demonstrated with western blotting. Rabbit anti-Spike RBD was used to detect the spike and S1 proteins, and mouse anti–Flag M2 to detect S2 protein (Supplementary materials).

### Intra-host diversity in infected hamsters

To examine intra-host diversity of the different SARS-CoV-2 strains, nasal washes were collected from infected hamsters at days 3 and 5 post-infection and whole genome sequencing conducted to determine the stability of the viral genome over the course of infection. As inferred by Illumina sequencing data, even minor genome variant populations (i.e. variants with frequency < 0.3) identified in the original inoculating strains **(Table 1)** remained present after 5 days of infection (**Table S2**). Interestingly, some genome variants, which were absent in the original strains, developed during hamster infection. The frequencies of these newly arisen variants were generally higher at 5 dpi than at 3 dpi (**Table S2**). This trend of increase implied that these new variants survive better in hamsters. Of note, there was no mixed population observed in HK-95, either in the original isolate or in animals after infection, which may indicate that the D614G variant has acquired a relatively stable genome constellation for replication in mammals; this stability in both human isolates and hamster infections also further supports the contention that hamsters are a suitable model for human SARS-CoV-2 infection.

## Discussion

SARS-CoV-2 continues to transmit globally and since its emergence in December 2019 has caused more than 20 million laboratory confirmed infections and over 0.7 million deaths, as of August 26 2020. Social distancing strategies have been adopted to slow down the dissemination of virus, in hope that effective vaccines and therapeutics will be developed and available soon enough to prevent further waves of transmission. More than seven months after transmission in humans was first observed, various genetic variants of SARS-CoV-2 have emerged and been analyzed but no clear evolutionary direction for SARS-CoV-2 is yet apparent (Deng et al., 2020; Fauver et al., 2020; Forster et al., 2020; Sanchez-Pacheco et al., 2020). Since the majority of people remain naive to this virus it is necessary to closely track any potential changes in the pathogenic properties of SARS-CoV-2 variants along the course of transmission. SARS-CoV-2 is of zoonotic origin and is expected to gain more adaptations as it circulates in humans. Attenuated variants with deletions in the spike protein S1/S2 junction and other regions have been detected in virus-infected cell cultures and in patient specimens (Lau et al., 2020b; Wong et al., 2020). One variant containing D614G in the spike protein has become the dominant population in many countries since March 2020 and is reported to infect cells more efficiently *in vitro* (Daniloski et al., 2020; Li et al., 2020; Zhang et al., 2020b). To understand if D614G variant virus may exhibit altered pathogenicity and transmissibility, we compared 5 SARS-CoV-2 isolates obtained from Hong Kong returnees from Wuhan, China, in February and travelers who had visited Europe and other countries after March 2020, using the hamster infection model. We found that the D614G bearing strain, HK-95, replicates more efficiently in the airways of infected hamsters in the lungs than strains lacking D614G (**Figure 2 and S2**). A contact transmission experiment showed that naive hamsters exposed to a HK-95-infected hamster have higher titers of virus in their airways, suggesting that the D614G variant is more highly transmissible than non-D614G SARS-CoV-2 strains (**Figure 5**).

Infection with SARS-CoV-2 can range from being asymptomatic to causing severe disease and fatalities; it is not yet well understood which viral factors, together with host factors and conditions, may determine pathogenicity in humans. Besides the insertion of a polybasic cleavage PRRA motif at the S1/S2 junction of the spike protein, which clearly contributes to increased virulence features, and was probably responsible for the initial SARS-CoV-2 cross species transmission event (Andersen et al., 2020; Lau et al., 2020b), other virulence elements are not well defined. The spike D614G variant, which was first recorded in January in China (hCoV-19/Zhejiang/HZ103/2020; 24 January 2020) and shortly thereafter in Germany (hCoV-19/Germany/BavPat1-Chvir929/2020; 28 January 2020), was found to have subsequently become the dominant population in multiple countries, suggesting SARS-CoV-2 is adapting to become more transmissible in humans. Host adaptation of an emerging virus in a naive population is generally believed to involve the virus gaining more efficient replicative ability while gradually decreasing its pathogenicity in the new host. It is not clear if the spike D614G variant will drive SARS-CoV-2 towards this evolutionary pathway. The more pathogenic features of the HK-95 strain, which carries the D614G substitution, observed in hamsters in this study could be due to the high replication efficiency of this strain in animals, given that hamsters are highly susceptible to SARS-CoV-2 infection. This is consistent with our transcriptome analysis of lung tissues from infected hamsters, which revealed no significant difference between host responses to infection with 614D or 614G strains (**Figure 4 and S4**). However, the contact transmission experiment demonstrated that HK-95 exhibits higher transmissibility to naive hamsters than HK-13 (**Figure 5**), despite these strains provoking similar pathogenic effects and transcriptome profiles in infected hamsters (**Figure 1, 2, 4 and S4**). If the spike D614G mutation is joined by other adaptative mutations as the virus further circulates in humans, it may be postulated that a variant with the D614G substitution together with a deletion in the polybasic cleavage site at the S1/S2 junction could arise, and if so, it may present as a much less pathogenic version of SARS-CoV-2 in the aftermath of the COVID-19 pandemic. Nevertheless, human interventions, such as mass vaccination as soon as vaccines are available, and preexisting immunity from prior infections are likely to drive the evolution of variants that can evade host immunity.

The increased transmissibility of the D614G SARS-CoV-2 strain is likely due to its higher replication ability. How the spike D614G mutation enhances virus replication and consequently transmissibility has not been defined. We and others have shown that D614G variant spike proteins are more efficiently cleaved into S1 and S2 subunits in cells (**Figure 5B**) (Daniloski et al., 2020; Zhang et al., 2020b). Two synonymous mutations, *5’UTR-C241T* and *Orf1a-C3037T*, and one nonsynonymous mutation, *Orf1b-P314L*, are consistently linked in D614G variant strains (**Table 1**). It remains to be investigated whether these mutations jointly contribute to the enhanced replication ability of the HK-95 SARS-CoV-2 isolate. Of note, *Orf1b-P314L* lies within the RNA-dependent RNA polymerase (RdRp) region, suggesting this mutation may play a causative role on enhanced viral replication. Further studies, including structural and functional analyses, will provide necessary information for understanding the molecular basis underlying the D614G-associated SARS-CoV-2 phenotype.

## Acknowledgements

The authors would like to thank Dr Jane Rayner for critical reading and editing of the manuscript. This study is partly supported by the Theme-Based Research Scheme (T11/707/15) and General Research Fund (17107019) of the Research Grants Council, Hong Kong Special Administrative Region, China, and the Sanming Project of Medicine in Shenzhen, China (No. SZSM201911014).

## Author contribution

B.W.M. and H.C. designed the studies; B.W.M., C.J.C., S-Y.L., S.D., P.C., A.J.Z., A.C-Y.L., H.L., S.L., T.T-L.N., H-Y.L., E.L-K.L., K.S-S.L,.P.W. and K-H.C. performance experiments;

B.W.M., C.J.C., S-Y.L., A.J.Z., K.K-W.T., J.F-W.C, K-Y.Y., G.K-H.S., and H.C. analyzed the data; B.W. M., C. J.C., G. K-H. S. and H.C. wrote the paper.

## Competing Interests statement

The authors declare no conflict of interests.

## Materials and Methods

### Virus

The SARS-CoV-2 isolates HK-8 (MT835139), HK-13 (MT835140), HK-15 (MT835141), HK-92 (MT835142) and HK-95 (MT835143) were isolated from specimens obtained from five laboratory-confirmed COVID-19 patients using Vero E6 cells (ATCC; CRL-15786). All experiments involving SARS-CoV-2 viruses were conducted in a Biosafety Level-3 laboratory. For animal challenge, viral stocks were prepared after two serial passages of isolated virus in Vero E6 cells in Dulbecco’s Modified Eagle Medium (DMEM) (Thermo Fisher Scientific) supplemented with 5% fetal bovine serum (Thermo Fisher Scientific), and 100 IU penicillin G/ml and 100 ml streptomycin sulfate/ml (Thermo Fisher Scientific). Virus titers were then determined by plaque assay using Vero E6 cells. Viral RNAs were also obtained from the supernatants of infected cells and then isolated using the QIAamp RNA Viral kit (Qiagen) and subjected to whole viral genome sequencing.

### Hamster infection

Female golden Syrian hamsters, aged 8-9 weeks old, were obtained from the Laboratory Animal Unit, the University of Hong Kong (HKU). All experiments were performed in a Biosafety Level-3 animal facility at the LKS Faculty of Medicine, HKU. All animal studies were approved by the Committee on the Use of Live Animals in Teaching and Research, HKU. Virus stocks were diluted with phosphate-buffered saline (PBS) to 2 x 10^4^ PFU/ml. Hamsters were anesthetized with ketamine (150mg/kg) and xylazine (10 mg/mg) and then intranasally inoculated with 50 ul of diluted virus stock containing 10^3^ PFU of virus or 50 ul PBS (mock infection control). Body weights were monitored daily for 5 days. Nasal washes were collected from hamsters at 3 and 5 dpi. Total nucleic acid was extracted from 140 ul of sample fluid using the QIAamp RNA Viral kit (Qiagen) and eluted with 30 ul of DEPC-treated water. Seven ul RNA was used for reverse transcription using MultiScribe Reverse Transcriptase (Thermofisher). cDNA was subsequently used for real-time qPCR using TB Green Premix Ex Taq (Tli RNase H Plus) (Takara). Viral RNA from nasal washes was also used for whole viral genome sequencing. Hamsters were euthanized at 5 dpi and lung tissues collected for histopathology, determination of viral load and RNA sequencing.

### Viral load determination and histopathology

Lung right lobes (superior, middle and inferior) were homogenized in 1 ml of PBS. After centrifugation at 12,000 rpm for 10 min, the supernatant was harvested, and viral titers determined by plaque assay using Vero E6 cells. Lung left superior lobes were fixed in 4 % paraformaldehyde and then processed for paraffin embedding. The 4 μm tissue sections were stained with haematoxylin and eosin for histopathological examination. Images were obtained with an Olympus BX53 semi-motorized fluorescence microscope using cellSens imaging software.

### RNA sequencing of hamster lung tissues

Lung left inferior lobes from hamsters were cut into pieces and lysed using the RA1 lysis buffer provided with the NucleoSpin*®* RNA Plus kit (Macherey-Nagel). RNA extraction was then performed according to the manufacturer’s instructions, including an on-column genomic DNA digestion step. 1 μg of high-quality total RNA (RIN>8) was used for cDNA library preparation with a KAPA mRNA HyperPrep Kit. The libraries were then denatured and diluted to optimal concentration, before being sequenced on an Illumina NovaSeq 6000 in a 151bp Paired-End format. RNA-seq data used in this study can be accessed in GEO under the accession number GSE156005.

### Transmission experiment

For each virus strain two hamsters were intranasally inoculated with 10^3^ PFU of virus. At twenty-four hours post-infection (1 dpi), each infected donor hamster was transferred to a new cage and co-housed with three naïve hamsters for 1 day. At 4 dpi, recipient hamsters were euthanized, and lung and nasal turbinate tissues collected for determination of viral load.

### Differential expression (DE) analysis

Sequencing reads were aligned to the merged golden hamster (Mesocricetus_auratus, V1.0, ENSEMBL v100) and the SARS-CoV-2 (NCBI Accession: NC_045512.2) genomes using STAR (Dobin et al., 2013). Read counts were extracted using the “--quantMode GeneCounts” argument with STAR for each sample. Counts were used to perform differential expression (DE) with DESeq2 (Love et al., 2014). DE thresholds required genes to have log2FoldChange ? 1 and a p-adjusted value ≤ 0.05 to be considered for downstream characterization of gene expression between conditions. Volcano plots were generated using the EnhancedVolcano R package (www.bioconductor.org/packages/release/bioc/vignettes/EnhancedVolcano/inst/doc/EnhancedVolcano.html). Normalized viral gene counts were extracted from DESeq2 output. Boxplots were generated with ggplot2 R package (Wickham, 2016). All analysis was performed through R 4.0 with custom R script.

### Principal component analysis

Log2FoldChange values for hamster DEGs for infections with each of the five isolates, contrasted to samples from uninfected animals, were extracted from the results of DESeq2. PCA analysis was performed on these values using the prcomp function from the stats R package. Scatterplots of PCA outputs were generated using the ggplot2 R package.

### Evolutionary analysis

The lack of annotation regarding mechanisms underlying the regulation of many biological networks in hamsters required the determination of a well-annotated close relative from which to infer the biological roles of differentially expressed genes in a species with a better representation of the dynamics of the SARS-CoV-2 host response in humans. Evolutionary analysis was performed in accordance to the protocol set by the author (Hall, 2013), using MEGA X software. Briefly, IL6 gene sequence homologs were acquired through an extensive search of the ENSEMBL (v100) database and their identity verified through nBLAST. Sequences were aligned using MUSCLE, with positions with less than 95% coverage being excluded (Edgar, 2004). The evolutionary history of IL6 was inferred using the Maximum Likelihood method and Tamura 3-parameter model (Tamura, 1992).

### Gene set enrichment analysis (GSEA)

Comprehensive lists of known mouse-hamster gene orthologues were compiled from the BioMart database (Smedley et al., 2015). Differentially expressed hamster genes which were identified to have a mouse orthologue form were retained from the orthologous gene list. These lists of orthologues were matched to genes categorized into Gene Ontology (GO) biological processes. The top 30 GO terms which were identified to have the most significant enrichment (FDR ≤ 0.05) across infected conditions were determined. A comparison between HK-95 and non-D614G isolates was also conducted. Heatmaps of significant GO groups were generated using the ComplexHeatmap R package (Gu et al., 2016).

### Whole viral genome sequencing

#### Reverse transcription and viral genome amplification using multiplex PCR

Viral RNA extracted from cell cultures and nasal washes was treated using the TURBO DNA-free Kit (ThermoFisher Scientific) to remove residual host DNA, followed by synthesis of single-strand cDNA using SuperScript IV reverse transcriptase (Invitrogen). The viral cDNA was then enriched through multiplex tiling polymerase chain reaction (PCR), as described in the ARTIC network (https://artic.network/ncov-2019) (Supplementary material).

#### Nanopore sequencing of viral genome

The input viral cDNA amplicons for individual samples were normalized to 5 ng, followed by end-repairing and adapter ligation according to official 1D sequencing protocols (SQK-LSK109, Oxford Nanopore). The libraries were sequenced on a Nanopore MinION device using an R9.4.1 flow cell for 48 hours. Nanopore sequencing data were analyzed using a modified Artic Network nCoV-2019 novel coronavirus bioinformatics protocol (Luo et al., 2020) (Supplementary material).

#### Illumina MiSeq sequencing of viral genome

A total of 100 ng of multiplex PCR amplicons were subjected to library preparation and dual-indexing using a KAPA HyperPrep Kit and a Unique Dual-Indexed Adapter Kit (Roche Applied Science) in accordance with the manufacturer’s instructions. Ligated libraries were then enriched by 6-cycle PCR amplification, followed by purification and size selection using AMPure XP beads (Beckman Coulter). The pooled libraries were sequenced using the MiSeq Reagent Kit V2 Nano on an Illumina MiSeq System. The Illumina MiSeq sequencing reads were then demultiplexed and mapped to the reference genome (accession number: NC_0.45512.2) using Samtools v1.7. Variants were called with Freebayes v1.0.0 (https://arxiv.org/abs/1207.3907) with the haploid setting, with a minimum base quality and depth of coverage of Q30 and 50x, respectively.

### Phylogenetic analysis

To determine the phylogenetic placement of our strains in the global phylogeny of SARS-CoV-2, a total of 100 SARS-CoV-2 genomes were downloaded from the GISAID severe acute respiratory syndrome coronavirus 2 data hub (Elbe and Buckland-Merrett, 2017). A phylogenetic tree was constructed with PhyML (v3.0) using the maximum likelihood algorithm. A best-fit substitution model for phylogenetic analysis was created using the Akaike information criterion, in which the general time reversible model with a fixed proportion of invariable sites (+I) was selected (Guindon et al., 2010). Bootstrap replicates were set at 1000×, and the maximum-likelihood phylogenetic tree was rooted on the earliest published genome of SARS-CoV-2 (accession no.: NC_045512.2).

## Notes

### Competing Interest Statement

The authors have declared no competing interest.

