## Supplementary materials for "SARS-CoV-2 spike D614G variant exhibits highly efficient replication and transmission in hamsters"

**Supplementary figures:**

**Figure S1.** Phylogenetic tree of SARS-CoV-2 isolates which contain either 614G (marked by bracket) or 614D (remaining viruses) in the spike protein. Isolates selected for use in this study are marked in red.

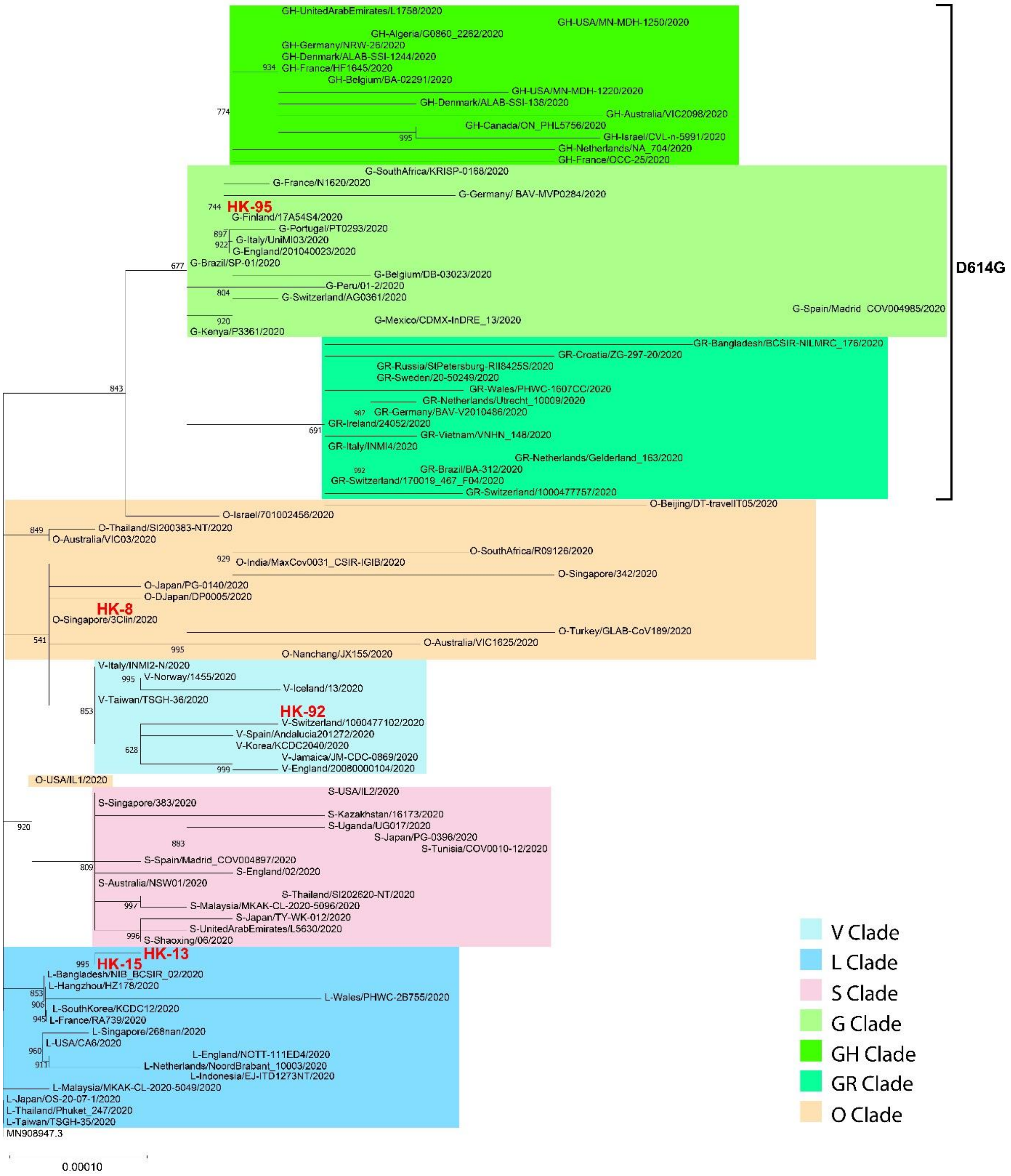

**Figure S2.** Real-time qPCR detection of viral RNA in nasal washes collected on day 5 post infection from hamsters intranasally challenged with different SARS-CoV-2 isolates.

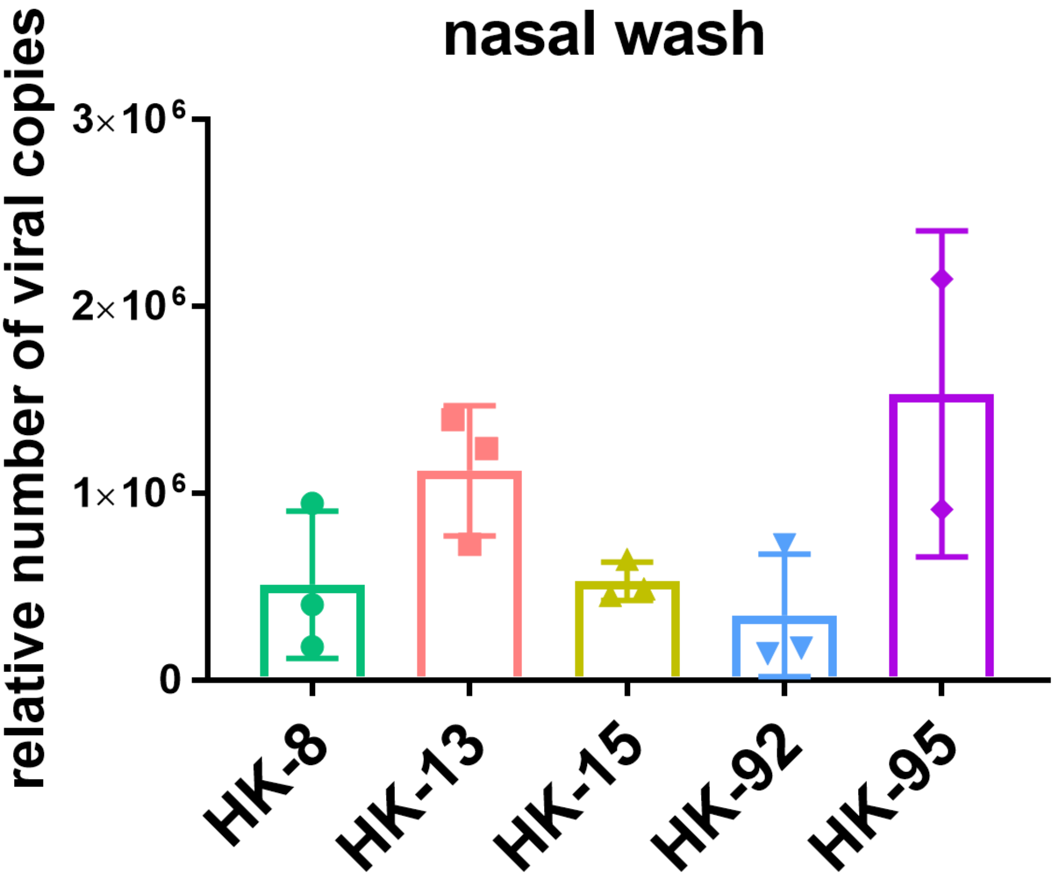

**Figure S3.** Cladogram describing the phylogenetics of the IL6 gene across taxa.

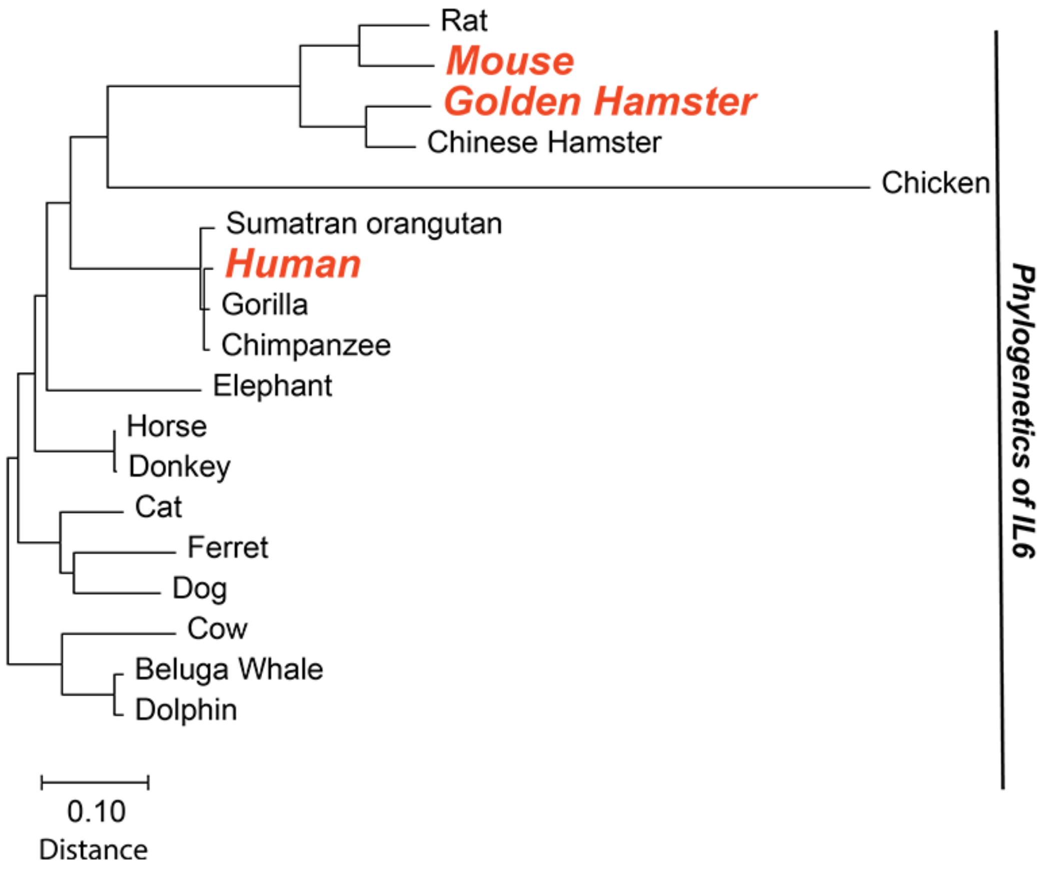

**Figure S4.** GO enrichment of differentially expressed genes in hamster lungs induced by various SARS-CoV-2 isolates compared to samples derived from hamsters infected with HK-95, at day 5 post-infection. Infection with HK-8 yielded significant GO enrichment in several categories relating to chromatin remodeling processes. GO enriched signals in response to infection with HK-8, compared to HK-95, identified several gene functional networks that are part of an enhanced host response to HK-8 SARS-CoV-2 infection over that induced by HK-95 infection.

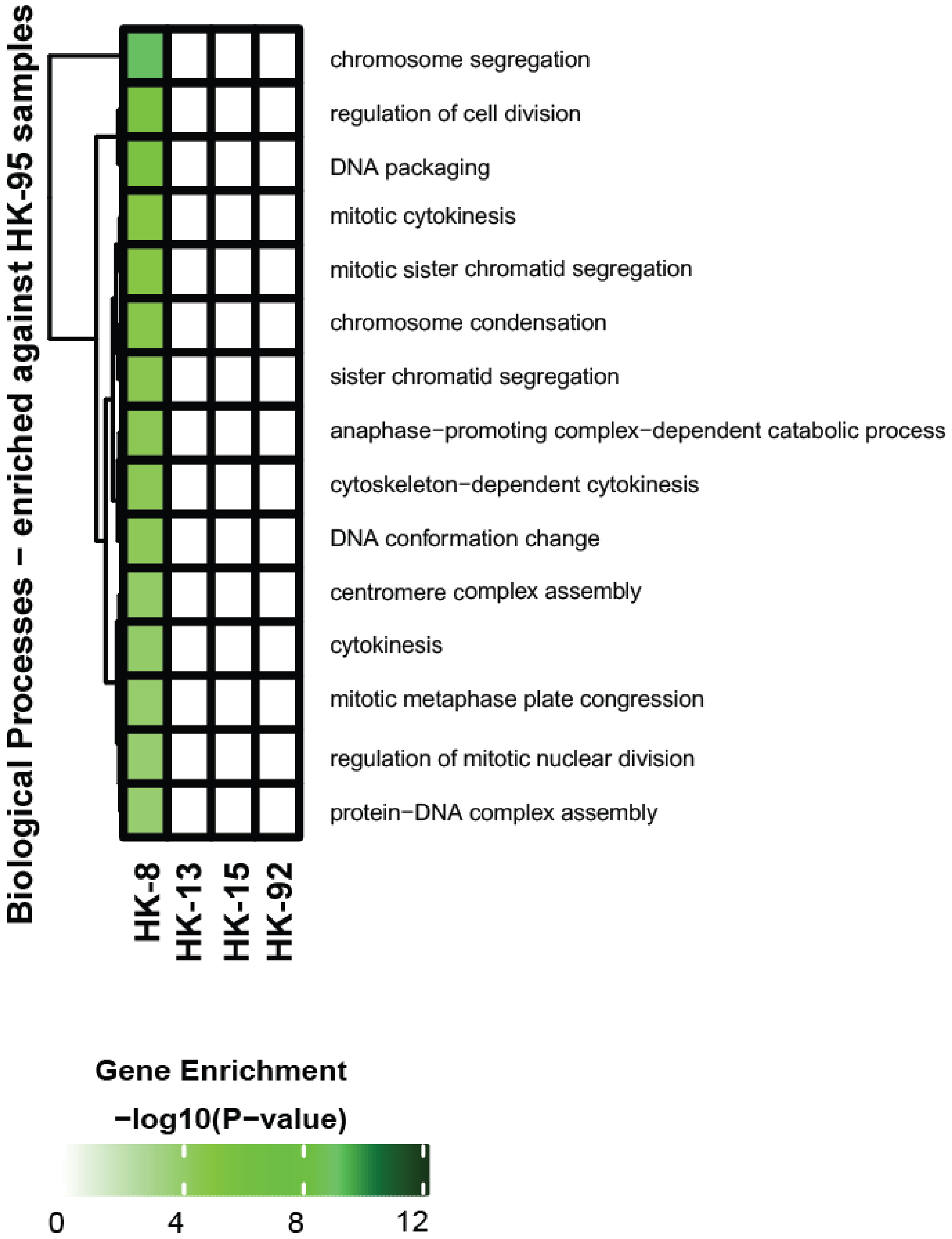

**Figure S5. Illustration of transmission experiment in hamsters.**

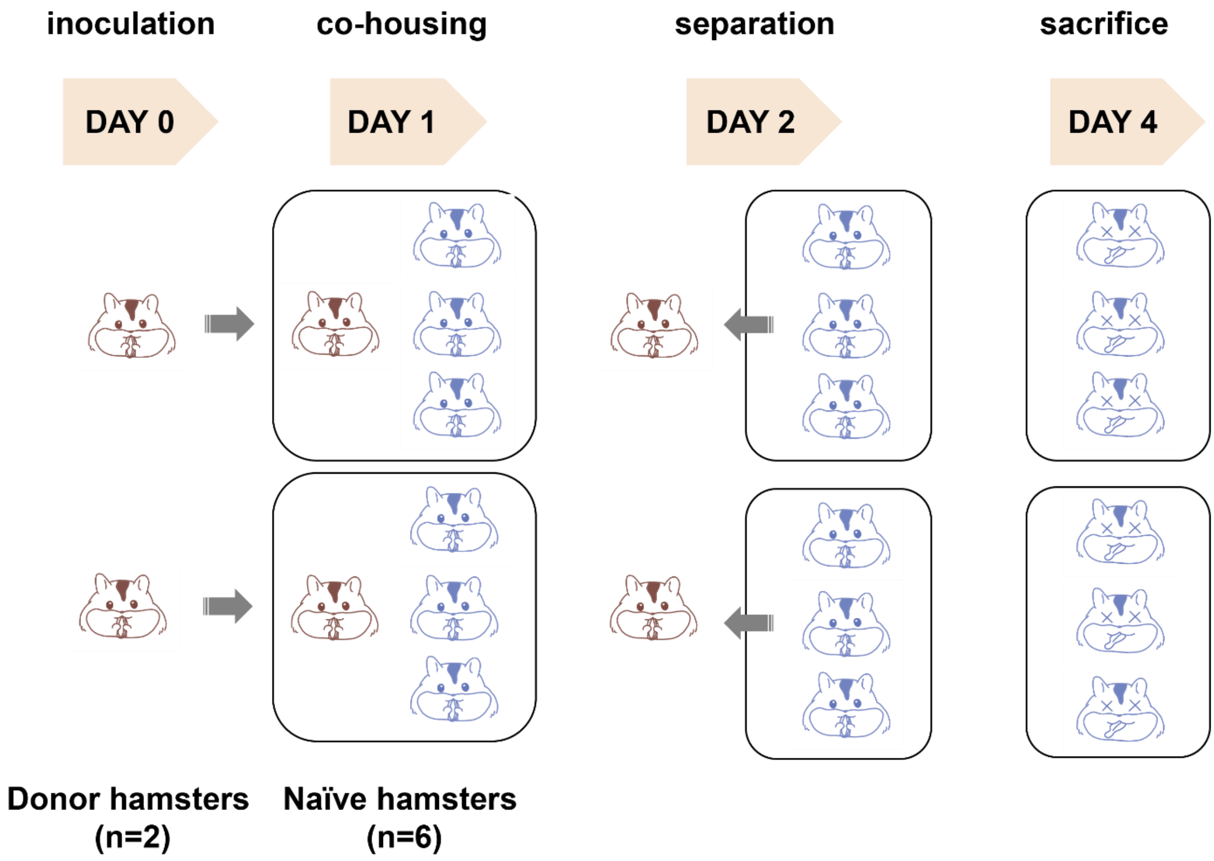

**Supplementary materials:**

### Multiplex PCR

The PCR contained a pool of 98 primer pairs, which generated overlapping 400-bp amplicons across the entire genome of SARS-CoV-2 (accession no: NC_045512) (primer sequences are shown in table S1). The final PCR mastermix (25μL) included 2.5μL of cDNA, 5.0μL of 5X Q5 Reaction Buffer, 0.5μL of 10mM dNTP mix, 0.25μL of Q5 Hot Start DNA Polymerase (New England Biolabs) and 3.6μL of 10μM primer pool 1 or 2, plus 13.15μL of nuclease-free water. The mixtures were incubated at 98°C for 30 s, followed by 35 cycles at 98°C for 15 s and 65°C for 5 min. The PCR amplicons were then purified using 1X Agencourt AMPure XP beads (Beckman Coulter).

### Bioinformatic analysis of Nanopore sequencing data

In brief, adapters were trimmed with Porechop v0.2.4, followed by primer sequence removal. The processed sequencing reads were then mapped to the SARS-CoV-2 isolate Wuhan-Hu-1 reference genome (accession number: NC_0.45512.2) using BWA v0.7.17 and Samtools v1.7. Variants were called with Medaka v0.11.5. The minimum depth of coverage and the minimum Phred score were set at 20 and 10, respectively [1]. The variants were then validated by mapping the sequencing reads to the reference genome with Minimap2 [2] and Clair v2 [3].

### Western blotting

For protein expression, plasmids containing the spike gene were transfected into HEK293T-ACE2 cells using TransIT-LT1 transfection reagent (Mirus). Cells were incubated for 48 hours and supernatants then boiled in 6x Laemmli sample buffer at 95˚C for 10 min. Samples were fractionated by 8% SDS-PAGE and then blotted onto nitrocellulose membranes (Bio-Rad). After blocking with 3% skim milk, membranes were incubated with rabbit anti-Spike RBD and mouse anti–Flag M2 (Sigma) at 1:1000 dilution. IRDye 680– or IRDye 780–labeled donkey anti-mouse or anti-rabbit secondary Abs (Li-Cor Biosciences) were used at a dilution of 1:5000. For visualization, membrane blots were scanned using the Odyssey imaging system (Li-Cor Biosciences).

1) Loman N, Rambaut A. nCoV-2019 novel coronavirus bioinformatics protocol. 2020. https://artic.network/ncov-2019/ncov2019-bioinformatics-sop.html (accessed Mar 10, 2020)
2) Li, H., Minimap2: pairwise alignment for nucleotide sequences. Bioinformatics, 34:3094-3100.
3) Ruibang Luo, Chak-Lim Wong, Yat-Sing Wong, Chi-Ian Tang, Chi-Man Liu, Chi-Ming Leung & Tak-Wah Lam, Exploring the limit of using a deep neural network on pileup data for germline variant calling.

**Table S1: Primers used in multiplex PCRs for SARS-CoV-2 genome amplification**

| Primer name | Sequence | Primer Start[a] | Primer end[a] | PCR Pool | Primer length | GC contents | Tm |
|---|---|---|---|---|---|---|---|
| nCoV-2019_1_LEFT | ACCAACCAACTTTCGATCTCTTGT | 30 | 54 | Pool 1 | 24 | 41.67 | 60.69 |
| nCoV-2019_1_RIGHT | CATCTTTAAGATGTTGACGTGCCTC | 385 | 410 | Pool 1 | 25 | 44 | 60.45 |
| nCoV-2019_2_LEFT | CTGTTTTACAGGTTCGCGACGT | 320 | 342 | Pool 2 | 22 | 50 | 61.67 |
| nCoV-2019_2_RIGHT | TAAGGATCAGTGCCAAGCTCGT | 704 | 726 | Pool 2 | 22 | 50 | 61.74 |
| nCoV-2019_3_LEFT | CGGTAATAAAGGAGCTGGTGGC | 642 | 664 | Pool 1 | 22 | 54.55 | 61.32 |
| nCoV-2019_3_RIGHT | AAGGTGTCTGCAATTCATAGCTCT | 1004 | 1028 | Pool 1 | 24 | 41.67 | 60.32 |
| nCoV-2019_4_LEFT | GGTGTATACTGCTGCCGTGAAC | 943 | 965 | Pool 2 | 22 | 54.55 | 61.56 |
| nCoV-2019_4_RIGHT | CACAAGTAGTGGCACCTTCTTTAGT | 1312 | 1337 | Pool 2 | 25 | 44 | 60.97 |
| nCoV-2019_5_LEFT | TGGTGAAACTTCATGGCAGACG | 1242 | 1264 | Pool 1 | 22 | 50 | 61.39 |
| nCoV-2019_5_RIGHT | ATTGATGTTGACTTTCTCTTTTTGGAGT | 1623 | 1651 | Pool 1 | 28 | 32.14 | 60.17 |
| nCoV-2019_6_LEFT | GGTGTTGTTGGAGAAGGTTCCG | 1573 | 1595 | Pool 2 | 22 | 54.55 | 61.64 |
| nCoV-2019_6_RIGHT | TAGCGGCCTTCTGTAAAACACG | 1942 | 1964 | Pool 2 | 22 | 50 | 61.18 |
| nCoV-2019_7_LEFT | ATCAGAGGCTGCTCGTGTTGTA | 1875 | 1897 | Pool 1 | 22 | 50 | 61.73 |
| nCoV-2019_7_RIGHT | TGCACAGGTGACAATTTGTCCA | 2247 | 2269 | Pool 1 | 22 | 45.45 | 60.95 |
| nCoV-2019_8_LEFT | AGAGTTTCTTAGAGACGGTTGGGA | 2181 | 2205 | Pool 2 | 24 | 45.83 | 61 |
| nCoV-2019_8_RIGHT | GCTTCAACAGCTTCACTAGTAGGT | 2568 | 2592 | Pool 2 | 24 | 45.83 | 60.56 |
| nCoV-2019_9_LEFT | TCCCACAGAAGTGTTAACAGAGGA | 2505 | 2529 | Pool 1 | 24 | 45.83 | 61.18 |
| nCoV-2019_9_RIGHT | ATGACAGCATCTGCCACAACAC | 2882 | 2904 | Pool 1 | 22 | 50 | 61.71 |
| nCoV-2019_10_LEFT | TGAGAAGTGCTCTGCCTATACAGT | 2826 | 2850 | Pool 2 | 24 | 45.83 | 61.12 |
| nCoV-2019_10_RIGHT | TCATCTAACCAATCTTCTTCTTGCTCT | 3183 | 3210 | Pool 2 | 27 | 37.04 | 60.31 |
| nCoV-2019_11_LEFT | GGAATTTGGTGCCACTTCTGCT | 3144 | 3166 | Pool 1 | 22 | 50 | 61.66 |
| nCoV-2019_11_RIGHT | TCATCAGATTCAACTTGCATGGCA | 3507 | 3531 | Pool 1 | 24 | 41.67 | 61.35 |
| nCoV-2019_12_LEFT | AAACATGGAGGAGGTGTTGCAG | 3460 | 3482 | Pool 2 | 22 | 50 | 61.08 |
| nCoV-2019_12_RIGHT | TTCACTCTTCATTTCCAAAAAGCTTGA | 3826 | 3853 | Pool 2 | 27 | 33.33 | 60.36 |
| nCoV-2019_13_LEFT | TCGCACAAATGTCTACTTAGCTGT | 3771 | 3795 | Pool 1 | 24 | 41.67 | 60.56 |
| nCoV-2019_13_RIGHT | ACCACAGCAGTTAAAACACCCT | 4142 | 4164 | Pool 1 | 22 | 45.45 | 60.36 |

| | | | | | | | |
|---|---|---|---|---|---|---|---|
| nCoV-2019_14_LEFT | CATCCAGATTCTGCCACTCTTGT | 4054 | 4077 | Pool 2 | 23 | 47.83 | 60.62 |
| nCoV-2019_14_RIGHT | AGTTTCCACACAGACAGGCATT | 4428 | 4450 | Pool 2 | 22 | 45.45 | 60.42 |
| nCoV-2019_15_LEFT | ACAGTGCTTAAAAAGTGTAAAAGTGCC | 4294 | 4321 | Pool 1 | 27 | 37.04 | 61.32 |
| nCoV-2019_15_RIGHT | AACAGAAACTGTAGCTGGCACT | 4674 | 4696 | Pool 1 | 22 | 45.45 | 60.16 |
| nCoV-2019_16_LEFT | AATTTGGAAGAAGCTGCTCGGT | 4636 | 4658 | Pool 2 | 22 | 45.45 | 60.82 |
| nCoV-2019_16_RIGHT | CACAACTTGCGTGTGGAGGTTA | 4995 | 5017 | Pool 2 | 22 | 50 | 61.32 |
| nCoV-2019_17_LEFT | CTTCTTTCTTTGAGAGAAGTGAGGACT | 4939 | 4966 | Pool 1 | 27 | 40.74 | 60.69 |
| nCoV-2019_17_RIGHT | TTTGTTGGAGTGTTAACAATGCAGT | 5296 | 5321 | Pool 1 | 25 | 36 | 60.11 |
| nCoV-2019_18_LEFT | TGGAAATACCCACAAGTTAATGGTTTAAC | 5230 | 5259 | Pool 2 | 29 | 34.48 | 60.69 |
| nCoV-2019_18_RIGHT | AGCTTGTTTACCACACGTACAAGG | 5620 | 5644 | Pool 2 | 24 | 45.83 | 61.51 |
| nCoV-2019_19_LEFT | GCTGTTATGTACATGGGCACACT | 5563 | 5586 | Pool 1 | 23 | 47.83 | 61.18 |
| nCoV-2019_19_RIGHT | TGTCCAACTTAGGGTCAATTTCTGT | 5932 | 5957 | Pool 1 | 25 | 40 | 60.4 |
| nCoV-2019_20_LEFT | ACAAAGAAAACAGTTACACAACAACCA | 5867 | 5894 | Pool 2 | 27 | 33.33 | 60.68 |
| nCoV-2019_20_RIGHT | ACGTGGCTTTATTAGTTGCATTGTT | 6247 | 6272 | Pool 2 | 25 | 36 | 60.28 |
| nCoV-2019_21_LEFT | TGGCTATTGATTATAAACACTACACACCC | 6167 | 6196 | Pool 1 | 29 | 37.93 | 61.49 |
| nCoV-2019_21_RIGHT | TAGATCTGTGTGGCCAACCTCT | 6528 | 6550 | Pool 1 | 22 | 50 | 60.83 |
| nCoV-2019_22_LEFT | ACTACCGAAGTTGTAGGAGACATTATACT | 6466 | 6495 | Pool 2 | 29 | 37.93 | 61.25 |
| nCoV-2019_22_RIGHT | ACAGTATTCTTTGCTATAGTAGTCGGC | 6846 | 6873 | Pool 2 | 27 | 40.74 | 60.73 |
| nCoV-2019_23_LEFT | ACAACTACTAACATAGTTACACGGTGT | 6718 | 6745 | Pool 1 | 27 | 37.04 | 60.26 |
| nCoV-2019_23_RIGHT | ACCAGTACAGTAGGTTGCAATAGTG | 7092 | 7117 | Pool 1 | 25 | 44 | 60.57 |
| nCoV-2019_24_LEFT | AGGCATGCCTTCTTACTGTACTG | 7035 | 7058 | Pool 2 | 23 | 47.83 | 60.37 |
| nCoV-2019_24_RIGHT | ACATTCTAACCATAGCTGAAATCGGG | 7389 | 7415 | Pool 2 | 26 | 42.31 | 61.19 |
| nCoV-2019_25_LEFT | GCAATTGTTTTTCAGCTATTTTGCAGT | 7305 | 7332 | Pool 1 | 27 | 33.33 | 60.73 |

| | | | | | | | |
|---|---|---|---|---|---|---|---|
| nCoV-2019_25_RIGHT | ACTGTAGTGACAAGTCTCTCGCA | 7671 | 7694 | Pool 1 | 23 | 47.83 | 61.3 |
| nCoV-2019_26_LEFT | TTGTGATACATTCTGTGCTGGTAGT | 7626 | 7651 | Pool 2 | 25 | 40 | 60.28 |
| nCoV-2019_26_RIGHT | TCCGCACTATCACCAACATCAG | 7997 | 8019 | Pool 2 | 22 | 50 | 60.42 |
| nCoV-2019_27_LEFT | ACTACAGTCAGCTTATGTGTCAACC | 7943 | 7968 | Pool 1 | 25 | 44 | 60.8 |
| nCoV-2019_27_RIGHT | AATACAAGCACCAAGGTCACGG | 8319 | 8341 | Pool 1 | 22 | 50 | 61.13 |
| nCoV-2019_28_LEFT | ACATAGAAGTTACTGGCGATAGTTGT | 8249 | 8275 | Pool 2 | 26 | 38.46 | 60.13 |
| nCoV-2019_28_RIGHT | TGTTTAGACATGACATGAACAGGTGT | 8635 | 8661 | Pool 2 | 26 | 38.46 | 60.91 |
| nCoV-2019_29_LEFT | ACTTGTGTTCCTTTTTGTTGCTGC | 8595 | 8619 | Pool 1 | 24 | 41.67 | 61.39 |
| nCoV-2019_29_RIGHT | AGTGTACTCTATAAGTTTTGATGGTGTGT | 8954 | 8983 | Pool 1 | 29 | 34.48 | 60.69 |
| nCoV-2019_30_LEFT | GCACAACTAATGGTGACTTTTTGCA | 8888 | 8913 | Pool 2 | 25 | 40 | 61.19 |
| nCoV-2019_30_RIGHT | ACCACTAGTAGATACACAAACACCAG | 9245 | 9271 | Pool 2 | 26 | 42.31 | 60.3 |
| nCoV-2019_31_LEFT | TTCTGAGTACTGTAGGCACGGC | 9204 | 9226 | Pool 1 | 22 | 54.55 | 62.03 |
| nCoV-2019_31_RIGHT | ACAGAATAAACACCAGGTAAGAATGAGT | 9557 | 9585 | Pool 1 | 28 | 35.71 | 60.69 |
| nCoV-2019_32_LEFT | TGGTGAATACAGTCATGTAGTTGCC | 9477 | 9502 | Pool 2 | 25 | 44 | 61.09 |
| nCoV-2019_32_RIGHT | AGCACATCACTACGCAACTTTAGA | 9834 | 9858 | Pool 2 | 24 | 41.67 | 60.56 |
| nCoV-2019_33_LEFT | ACTTTTGAAGAAGCTGCGCTGT | 9784 | 9806 | Pool 1 | 22 | 45.45 | 61.58 |
| nCoV-2019_33_RIGHT | TGGACAGTAAACTACGTCATCAAGC | 10146 | 10171 | Pool 1 | 25 | 44 | 61.08 |
| nCoV-2019_34_LEFT | TCCCATCTGGTAAAGTTGAGGGT | 10076 | 10099 | Pool 2 | 23 | 47.83 | 61.02 |
| nCoV-2019_34_RIGHT | AGTGAAATTGGGCCTCATAGCA | 10437 | 10459 | Pool 2 | 22 | 45.45 | 60.03 |
| nCoV-2019_35_LEFT | TGTTCGCATTCAACCAGGACAG | 10362 | 10384 | Pool 1 | 22 | 50 | 61.39 |
| nCoV-2019_35_RIGHT | ACTTCATAGCCACAAGGTTAAAGTCA | 10737 | 10763 | Pool 1 | 26 | 38.46 | 60.69 |
| nCoV-2019_36_LEFT | TTAGCTTGGTTGTACGCTGCTG | 10666 | 10688 | Pool 2 | 22 | 50 | 61.44 |
| nCoV-2019_36_RIGHT | GAACAAAGACCATTGAGTACTCTGGA | 11048 | 11074 | Pool 2 | 26 | 42.31 | 60.74 |
| nCoV-2019_37_LEFT | ACACACCACTGGTTGTTACTCAC | 10999 | 11022 | Pool 1 | 23 | 47.83 | 60.93 |

| | | | | | | | |
|---|---|---|---|---|---|---|---|
| nCoV-2019_37_RIGHT | GTCCACACTCTCCTAGCA CCAT | 11372 | 11394 | Pool 1 | 22 | 54.55 | 61.48 |
| nCoV-2019_38_LEFT | ACTGTGTTATGTATGCAT CAGCTGT | 11306 | 11331 | Pool 2 | 25 | 40 | 60.86 |
| nCoV-2019_38_RIGHT | CACCAAGAGTCAGTCTAA AGTAGCG | 11668 | 11693 | Pool 2 | 25 | 48 | 61.13 |
| nCoV-2019_39_LEFT | AGTATTGCCCTATTTTCTT CATAACTGGT | 11555 | 11584 | Pool 1 | 29 | 34.48 | 61 |
| nCoV-2019_39_RIGHT | TGTAACTGGACACATTGA GCCC | 11927 | 11949 | Pool 1 | 22 | 50 | 60.55 |
| nCoV-2019_40_LEFT | TGCACATCAGTAGTCTTA CTCTCAGT | 11863 | 11889 | Pool 2 | 26 | 42.31 | 61.25 |
| nCoV-2019_40_RIGHT | CATGGCTGCATCACGGTC AAAT | 12234 | 12256 | Pool 2 | 22 | 50 | 62.09 |
| nCoV-2019_41_LEFT | GTTCCCTTCCATCATATG CAGCT | 12110 | 12133 | Pool 1 | 23 | 47.83 | 60.75 |
| nCoV-2019_41_RIGHT | TGGTATGACAACCATTAG TTTGGCT | 12465 | 12490 | Pool 1 | 25 | 40 | 60.75 |
| nCoV-2019_42_LEFT | TGCAAGAGATGGTTGTGT TCCC | 12417 | 12439 | Pool 2 | 22 | 50 | 61.08 |
| nCoV-2019_42_RIGHT | CCTACCTCCCTTTGTTGTG TTGT | 12779 | 12802 | Pool 2 | 23 | 47.83 | 60.69 |
| nCoV-2019_43_LEFT | TACGACAGATGTCTTGTG CTGC | 12710 | 12732 | Pool 1 | 22 | 50 | 60.93 |
| nCoV-2019_43_RIGHT | AGCAGCATCTACAGCAA AAGCA | 13074 | 13096 | Pool 1 | 22 | 45.45 | 61.14 |
| nCoV-2019_44_LEFT | TGCCACAGTACGTCTACA AGCT | 13005 | 13027 | Pool 2 | 22 | 50 | 61.66 |
| nCoV-2019_44_RIGHT | AACCTTTCCACATACCGC AGAC | 13378 | 13400 | Pool 2 | 22 | 50 | 60.87 |
| nCoV-2019_45_LEFT | TACCTACAACTTGTGCTA ATGACCC | 13319 | 13344 | Pool 1 | 25 | 44 | 60.57 |
| nCoV-2019_45_RIGHT | AAATTGTTTCTTCATGTTG GTAGTTAGAGA | 13669 | 13699 | Pool 1 | 30 | 30 | 60.01 |
| nCoV-2019_46_LEFT | TGTCGCTTCCAAGAAAAG GACG | 13599 | 13621 | Pool 2 | 22 | 50 | 61.38 |
| nCoV-2019_46_RIGHT | CACGTTCACCTAAGTTGG CGTA | 13962 | 13984 | Pool 2 | 22 | 50 | 60.86 |
| nCoV-2019_47_LEFT | AGGACTGGTATGATTTTG TAGAAAACCC | 13918 | 13946 | Pool 1 | 28 | 39.29 | 61.42 |
| nCoV-2019_47_RIGHT | AATAACGGTCAAAGAGTT TTAACCTCTC | 14271 | 14299 | Pool 1 | 28 | 35.71 | 60.06 |
| nCoV-2019_48_LEFT | TGTTGACACTGACTTAAC AAAGCCT | 14207 | 14232 | Pool 2 | 25 | 40 | 61.09 |
| nCoV-2019_48_RIGHT | TAGATTACCAGAAGCAGC GTGC | 14579 | 14601 | Pool 2 | 22 | 50 | 60.74 |
| nCoV-2019_49_LEFT | AGGAATTACTTGTGTATG CTGCTGA | 14545 | 14570 | Pool 1 | 25 | 40 | 60.57 |

| | | | | | | | |
|---|---|---|---|---|---|---|---|
| nCoV-2019_49_RIGHT | TGACGATGACTTGGTTAG CATTAATACA | 14898 | 14926 | Pool 1 | 28 | 35.71 | 61.05 |
| nCoV-2019_50_LEFT | GTTGATAAGTACTTTGAT TGTTACGATGGT | 14865 | 14895 | Pool 2 | 30 | 33.33 | 60.59 |
| nCoV-2019_50_RIGHT | TAACATGTTGTGCCAACC ACCA | 15224 | 15246 | Pool 2 | 22 | 45.45 | 60.95 |
| nCoV-2019_51_LEFT | TCAATAGCCGCCACTAGA GGAG | 15171 | 15193 | Pool 1 | 22 | 54.55 | 61.34 |
| nCoV-2019_51_RIGHT | AGTGCATTAACATTGGCC GTGA | 15538 | 15560 | Pool 1 | 22 | 45.45 | 61.14 |
| nCoV-2019_52_LEFT | CATCAGGAGATGCCACA ACTGC | 15481 | 15503 | Pool 2 | 22 | 54.55 | 61.83 |
| nCoV-2019_52_RIGHT | GTTGAGAGCAAAATTCAT GAGGTCC | 15861 | 15886 | Pool 2 | 25 | 44 | 60.62 |
| nCoV-2019_53_LEFT | AGCAAAATGTTGGACTGA GACTGA | 15827 | 15851 | Pool 1 | 24 | 41.67 | 60.69 |
| nCoV-2019_53_RIGHT | AGCCTCATAAAACTCAGG TTCCC | 16186 | 16209 | Pool 1 | 23 | 47.83 | 60.31 |
| nCoV-2019_54_LEFT | TGAGTTAACAGGACACAT GTTAGACA | 16118 | 16144 | Pool 2 | 26 | 38.46 | 60.18 |
| nCoV-2019_54_RIGHT | AACCAAAAACTTGTCCAT TAGCACA | 16485 | 16510 | Pool 2 | 25 | 36 | 60.11 |
| nCoV-2019_55_LEFT | ACTCAACTTTACTTAGGA GGTATGAGCT | 16416 | 16444 | Pool 1 | 28 | 39.29 | 61.43 |
| nCoV-2019_55_RIGHT | GGTGTACTCTCCTATTTG TACTTTACTGT | 16804 | 16833 | Pool 1 | 29 | 37.93 | 60.54 |
| nCoV-2019_56_LEFT | ACCTAGACCACCACTTAA CCGA | 16748 | 16770 | Pool 2 | 22 | 50 | 60.49 |
| nCoV-2019_56_RIGHT | ACACTATGCGAGCAGAA GGGTA | 17130 | 17152 | Pool 2 | 22 | 50 | 61.21 |
| nCoV-2019_57_LEFT | ATTCTACACTCCAGGGAC CACC | 17065 | 17087 | Pool 1 | 22 | 54.55 | 61.16 |
| nCoV-2019_57_RIGHT | GTAATTGAGCAGGGTCGC CAAT | 17430 | 17452 | Pool 1 | 22 | 50 | 61.26 |
| nCoV-2019_58_LEFT | TGATTTGAGTGTTGTCAA TGCCAGA | 17381 | 17406 | Pool 2 | 25 | 40 | 61.44 |
| nCoV-2019_58_RIGHT | CTTTTCTCCAAGCAGGGT TACGT | 17738 | 17761 | Pool 2 | 23 | 47.83 | 61.06 |
| nCoV-2019_59_LEFT | TCACGCATGATGTTTCAT CTGCA | 17674 | 17697 | Pool 1 | 23 | 43.48 | 61.42 |
| nCoV-2019_59_RIGHT | AAGAGTCCTGTTACATTT TCAGCTTG | 18036 | 18062 | Pool 1 | 26 | 38.46 | 60.02 |
| nCoV-2019_60_LEFT | TGATAGAGACCTTTATGA CAAGTTGCA | 17966 | 17993 | Pool 2 | 27 | 37.04 | 60.53 |
| nCoV-2019_60_RIGHT | GGTACCAACAGCTTCTCT AGTAGC | 18324 | 18348 | Pool 2 | 24 | 50 | 60.44 |
| nCoV-2019_61_LEFT | TGTTTATCACCCGCGAAG AAGC | 18253 | 18275 | Pool 1 | 22 | 50 | 61.5 |

| | | | | | | | |
|---|---|---|---|---|---|---|---|
| nCoV-2019_61_RIGHT | ATCACATAGACAACAGGTGCGC | 18650 | 18672 | Pool 1 | 22 | 50 | 61.25 |
| nCoV-2019_62_LEFT | GGCACATGGCTTTGAGTTGACA | 18596 | 18618 | Pool 2 | 22 | 50 | 61.91 |
| nCoV-2019_62_RIGHT | GTTGAACCTTTCTACAAGCCGC | 18957 | 18979 | Pool 2 | 22 | 50 | 60.35 |
| nCoV-2019_63_LEFT | TGTTAAGCGTGTTGACTGGACT | 18896 | 18918 | Pool 1 | 22 | 45.45 | 60.16 |
| nCoV-2019_63_RIGHT | ACAAACTGCCACCATCACAACC | 19275 | 19297 | Pool 1 | 22 | 50 | 61.85 |
| nCoV-2019_64_LEFT | TCGATAGATATCCTGCTAATTCCATTGT | 19204 | 19232 | Pool 2 | 28 | 35.71 | 60.11 |
| nCoV-2019_64_RIGHT | AGTCTTGTAAAAGTGTTCCAGAGGT | 19591 | 19616 | Pool 2 | 25 | 40 | 60.1 |
| nCoV-2019_65_LEFT | GCTGGCTTTAGCTTGTGGGTTT | 19548 | 19570 | Pool 1 | 22 | 50 | 61.92 |
| nCoV-2019_65_RIGHT | TGTCAGTCATAGAACAAACACCAATAGT | 19911 | 19939 | Pool 1 | 28 | 35.71 | 60.9 |
| nCoV-2019_66_LEFT | GGGTGTGGACATTGCTGCTAAT | 19844 | 19866 | Pool 2 | 22 | 50 | 61.21 |
| nCoV-2019_66_RIGHT | TCAATTTCCATTTGACTCCTGGGT | 20231 | 20255 | Pool 2 | 24 | 41.67 | 60.45 |
| nCoV-2019_67_LEFT | GTTGTCCAACAATTACCTGAAACTTACT | 20172 | 20200 | Pool 1 | 28 | 35.71 | 60.43 |
| nCoV-2019_67_RIGHT | CAACCTTAGAAACTACAGATAAATCTTGGG | 20542 | 20572 | Pool 1 | 30 | 36.67 | 60.4 |
| nCoV-2019_68_LEFT | ACAGGTTCATCTAAGTGTGTGTGT | 20472 | 20496 | Pool 2 | 24 | 41.67 | 60.14 |
| nCoV-2019_68_RIGHT | CTCCTTTATCAGAACCAGCACCA | 20867 | 20890 | Pool 2 | 23 | 47.83 | 60.31 |
| nCoV-2019_69_LEFT | TGTCGCAAAATATACTCAACTGTGTCA | 20786 | 20813 | Pool 1 | 27 | 37.04 | 61.43 |
| nCoV-2019_69_RIGHT | TCTTTATAGCCACGGAACCTCCA | 21146 | 21169 | Pool 1 | 23 | 47.83 | 61.14 |
| nCoV-2019_70_LEFT | ACAAAAGAAAATGACTCTAAAGAGGGTTT | 21075 | 21104 | Pool 2 | 29 | 31.03 | 60.13 |
| nCoV-2019_70_RIGHT | TGACCTTCTTTTAAAGACATAACAGCAG | 21427 | 21455 | Pool 2 | 28 | 35.71 | 60.27 |
| nCoV-2019_71_LEFT | ACAAATCCAATTCAGTTGTCTTCCTATTC | 21357 | 31386 | Pool 1 | 29 | 34.48 | 60.54 |
| nCoV-2019_71_RIGHT | TGGAAAAGAAAGGTAAGAACAAGTCCT | 21716 | 21743 | Pool 1 | 27 | 37.04 | 60.8 |
| nCoV-2019_72_LEFT | ACACGTGGTGTTTATTACCCTGAC | 21658 | 21682 | Pool 2 | 24 | 45.83 | 61.04 |
| nCoV-2019_72_RIGHT | ACTCTGAACTCACTTTCCATCCAAC | 22013 | 22038 | Pool 2 | 25 | 44 | 60.97 |

| | | | | | | | |
|---|---|---|---|---|---|---|---|
| nCoV-2019_73_LEFT | CAATTTTGTAATGATCCATTTTTGGGTGT | 21961 | 21990 | Pool 1 | 29 | 31.03 | 60.29 |
| nCoV-2019_73_RIGHT | CACCAGCTGTCCAACCTGAAGA | 22324 | 22346 | Pool 1 | 22 | 54.55 | 62.45 |
| nCoV-2019_74_LEFT | ACATCACTAGGTTTCAAACTTTACTTGC | 22262 | 22290 | Pool 2 | 28 | 35.71 | 60.68 |
| nCoV-2019_74_RIGHT | GCAACACAGTTGCTGATTCTCTTC | 22626 | 22650 | Pool 2 | 24 | 45.83 | 60.85 |
| nCoV-2019_75_LEFT | AGAGTCCAACCAACAGAATCTATTGT | 22516 | 22542 | Pool 1 | 26 | 38.46 | 60.24 |
| nCoV-2019_75_RIGHT | ACCACCAACCTTAGAATCAAGATTGT | 22877 | 22903 | Pool 1 | 26 | 38.46 | 60.69 |
| nCoV-2019_76_LEFT | AGGGCAAACTGGAAAGATTGCT | 22797 | 22819 | Pool 2 | 22 | 45.45 | 60.76 |
| nCoV-2019_76_RIGHT | ACACCTGTGCCTGTTAAACCAT | 23192 | 23214 | Pool 2 | 22 | 45.45 | 60.42 |
| nCoV-2019_77_LEFT | CCAGCAACTGTTTGTGGACCTA | 23122 | 23144 | Pool 1 | 22 | 50 | 60.75 |
| nCoV-2019_77_RIGHT | CAGCCCCTATTAAACAGCCTGC | 23500 | 23522 | Pool 1 | 22 | 54.55 | 61.59 |
| nCoV-2019_78_LEFT | CAACTTACTCCTACTTGGCGTGT | 23443 | 23466 | Pool 2 | 23 | 47.83 | 60.55 |
| nCoV-2019_78_RIGHT | TGTGTACAAAAACTGCCATATTGCA | 23822 | 23847 | Pool 2 | 25 | 36 | 60.22 |
| nCoV-2019_79_LEFT | GTGGTGATTCAACTGAATGCAGC | 23789 | 23812 | Pool 1 | 23 | 47.83 | 60.92 |
| nCoV-2019_79_RIGHT | CATTTCATCTGTGAGCAAAGGTGG | 24145 | 24169 | Pool 1 | 24 | 45.83 | 60.62 |
| nCoV-2019_80_LEFT | TTGCCTTGGTGATATTGCTGCT | 24078 | 24100 | Pool 2 | 22 | 45.45 | 60.89 |
| nCoV-2019_80_RIGHT | TGGAGCTAAGTTGTTTAACAAGCG | 24443 | 24467 | Pool 2 | 24 | 41.67 | 60.02 |
| nCoV-2019_81_LEFT | GCACTTGGAAAACTTCAAGATGTGG | 24391 | 24416 | Pool 1 | 25 | 44 | 61.24 |
| nCoV-2019_81_RIGHT | GTGAAGTTCTTTTCTTGTGCAGGG | 24765 | 24789 | Pool 1 | 24 | 45.83 | 60.73 |
| nCoV-2019_82_LEFT | GGGCTATCATCTTATGTCCTTCCCT | 24696 | 24721 | Pool 2 | 25 | 48 | 61.52 |
| nCoV-2019_82_RIGHT | TGCCAGAGATGTCACCTAAATCAA | 25052 | 25076 | Pool 2 | 24 | 41.67 | 60.02 |
| nCoV-2019_83_LEFT | TCCTTTGCAACCTGAATTAGACTCA | 24978 | 25003 | Pool 1 | 25 | 40 | 60.46 |
| nCoV-2019_83_RIGHT | TTTGACTCCTTTGAGCACTGGC | 25347 | 25369 | Pool 1 | 22 | 50 | 61.33 |
| nCoV-2019_84_LEFT | TGCTGTAGTTGTCTCAAGGGCT | 25279 | 25301 | Pool 2 | 22 | 50 | 61.61 |
| nCoV-2019_84_RIGHT | AGGTGTGAGTAAACTGTTACAAACAAC | 25646 | 25673 | Pool 2 | 27 | 37.04 | 60.36 |

| | | | | | | | |
|---|---|---|---|---|---|---|---|
| nCoV-2019_85_LEFT | ACTAGCACTCTCCAAGGGTGTT | 25601 | 25623 | Pool 1 | 22 | 50 | 61.03 |
| nCoV-2019_85_RIGHT | ACACAGTCTTTTACTCCAGATTCCC | 25969 | 25994 | Pool 1 | 25 | 44 | 60.51 |
| nCoV-2019_86_LEFT | TCAGGTGATGGCACAACAAGTC | 25902 | 25924 | Pool 2 | 22 | 50 | 61.07 |
| nCoV-2019_86_RIGHT | ACGAAAGCAAGAAAAAGAAGTACGC | 26290 | 26315 | Pool 2 | 25 | 40 | 61.01 |
| nCoV-2019_87_LEFT | CGACTACTAGCGTGCCTTTGTA | 26197 | 26219 | Pool 1 | 22 | 50 | 60.16 |
| nCoV-2019_87_RIGHT | ACTAGGTTCCATTGTTCAAGGAGC | 26566 | 26590 | Pool 1 | 24 | 45.83 | 60.81 |
| nCoV-2019_88_LEFT | CCATGGCAGATTCCAACGGTAC | 26520 | 26542 | Pool 2 | 22 | 54.55 | 61.58 |
| nCoV-2019_88_RIGHT | TGGTCAGAATAGTGCCATGGAGT | 26890 | 26913 | Pool 2 | 23 | 47.83 | 61.4 |
| nCoV-2019_89_LEFT | GTACGCGTTCCATGTGGTCATT | 26835 | 26857 | Pool 1 | 22 | 50 | 61.5 |
| nCoV-2019_89_RIGHT | ACCTGAAAGTCAACGAGATGAAACA | 27202 | 27227 | Pool 1 | 25 | 40 | 60.91 |
| nCoV-2019_90_LEFT | ACACAGACCATTCCAGTAGCAGT | 27141 | 27164 | Pool 2 | 23 | 47.83 | 61.58 |
| nCoV-2019_90_RIGHT | TGAAATGGTGAATTGCCCTCGT | 27511 | 27533 | Pool 2 | 22 | 45.45 | 60.82 |
| nCoV-2019_91_LEFT | TCACTACCAAGAGTGTGTTAGAGGT | 27446 | 27471 | Pool 1 | 25 | 44 | 60.93 |
| nCoV-2019_91_RIGHT | TTCAAGTGAGAACCAAAAGATAATAAGCA | 27825 | 27854 | Pool 1 | 29 | 31.03 | 60.03 |
| nCoV-2019_92_LEFT | TTTGTGCTTTTTAGCCTTTCTGCT | 27784 | 27808 | Pool 2 | 24 | 37.5 | 60.14 |
| nCoV-2019_92_RIGHT | AGGTTCCTGGCAATTAATTGTAAAAGG | 28145 | 28172 | Pool 2 | 27 | 37.04 | 60.53 |
| nCoV-2019_93_LEFT | TGAGGCTGGTTCTAAATCACCCA | 28081 | 28104 | Pool 1 | 23 | 47.83 | 61.59 |
| nCoV-2019_93_RIGHT | AGGTCTTCCTTGCCATGTTGAG | 28442 | 28464 | Pool 1 | 22 | 50 | 60.55 |
| nCoV-2019_94_LEFT | GGCCCCAAGGTTTACCCAATAA | 28394 | 28416 | Pool 2 | 22 | 50 | 60.56 |
| nCoV-2019_94_RIGHT | TTTGGCAATGTTGTTCCTTGAGG | 28756 | 28779 | Pool 2 | 23 | 43.48 | 60.18 |
| nCoV-2019_95_LEFT | TGAGGGAGCCTTGAATACACCA | 28677 | 28699 | Pool 1 | 22 | 50 | 61.1 |
| nCoV-2019_95_RIGHT | CAGTACGTTTTTGCCGAGGCTT | 29041 | 29063 | Pool 1 | 22 | 50 | 61.95 |
| nCoV-2019_96_LEFT | GCCAACAACAACAAGGCCAAAC | 28985 | 29007 | Pool 2 | 22 | 50 | 61.82 |
| nCoV-2019_96_RIGHT | TAGGCTCTGTTGGTGGGAATGT | 29356 | 29378 | Pool 2 | 22 | 50 | 61.36 |
| nCoV-2019_97_LEFT | TGGATGACAAAGATCCAAATTTCAAAGA | 29288 | 29316 | Pool 1 | 28 | 32.14 | 60.22 |

| | | | | | | | |
|---|---|---|---|---|---|---|---|
| nCoV-2019_97_RIGHT | ACACACTGATTAAAGATTGCTATGTGAG | 29665 | 29693 | Pool 1 | 28 | 35.71 | 60.17 |
| nCoV-2019_98_LEFT | AACAATTGCAACAATCCATGAGCA | 29486 | 29510 | Pool 2 | 24 | 37.5 | 60.5 |
| nCoV-2019_98_RIGHT | TTCTCCTAAGAAGCTATTAAAATCACATGG | 29836 | 29866 | Pool 2 | 30 | 33.33 | 60.01 |

[a] Nucleotide positions are based on Wuhan-Hu-1 (accession no. NC_045512.2)

**Supplementary Table 2: Genome variants (and their frequencies) identified in SARS-CoV-2 strains after infection in cell culture and hamster infection model**

| Isolate | Infection model | Days post-infection | Genome variant (Variant frequency)[#] | | | | | | | | |
|---|---|---|---|---|---|---|---|---|---|---|---|
| HK-8 | Cell culture | -- | 5310 C>T, Orf1a T1682I (VF=0.27) * | 8092 C>T, Orf1a L2609L (VF=1) | 11083 G>T, Orf1a L3606F (VF=1) | 21137 A>G, Orf1b K2557R (VF=0.07)** | NIL | | | | |
| | Hamster# 8.1 | 3 days | 5310 C>T, Orf1a T1682I (VF=0.41) * | 8092 C>T, Orf1a L2609L (VF=1) | 11083 G>T, Orf1a L3606F (VF=1) | 21137 A>G, Orf1b K2557R (VF=0.09)** | 29747 G>T, 3' Untranslated region (3'UTR) (VF=0.03)**[¶] | | | | |
| | | 5 days | 5310 C>T, Orf1a T1682I (VF=0.53) * | 8092 C>T, Orf1a L2609L (VF=1) | 11083 G>T, Orf1a L3606F (VF=1) | 21137 A>G, Orf1b K2557R (VF= 0.1) ** | 29747 G>T, 3' Untranslated region (3'UTR) (VF=0.02)**[¶] | | | | |
| | Hamster# 8.2 | 3 days | 5310 C>T, Orf1a T1682I (VF=0.24) * | 8092 C>T, Orf1a L2609L (VF=1) | 11083 G>T, Orf1a L3606F (VF=1) | 21137 A>G, Orf1b K2557R (VF=0.09) ** | 29747 G>T, 3' Untranslated region (3'UTR) (VF=0.03)**[¶] | | | | |
| | | 5 days | 5310 C>T, Orf1a T1682I (VF=0.45) * | 8092 C>T, Orf1a L2609L (VF=1) | 11083 G>T, Orf1a L3606F (VF=1) | 21137 A>G, Orf1b K2557R (VF=0.07) ** | 29747 G>T, 3' Untranslated region (3'UTR) (VF=0.10)**[¶] | | | | |
| | Hamster# 8.3 | 3 days | 5310 C>T, Orf1a T1682I (VF=0.37) * | 8092 C>T, Orf1a L2609L (VF=1) | 11083 G>T, Orf1a L3606F (VF=1) | 21137 A>G, Orf1b K2557R (VF=0.11) * | 29747 G>T, 3' Untranslated region (3'UTR) (VF=0.07)**[¶] | | | | |
| | | 5 days | 5310 C>T, Orf1a T1682I (VF= 0.39) * | 8092 C>T, Orf1a L2609L (VF=1) | 11083 G>T, Orf1a L3606F (VF=1) | 21137 A>G, Orf1b K2557R (VF=0.09) ** | 29747 G>T, 3' Untranslated region (3'UTR) (VF=0.08)**[¶] | | | | |
| HK-13 | Cell culture | -- | 17423 A>G, Orf1b Y1319C (VF=1) | 20667 A>G, Orf1b Q2400Q (VF=0.99) | NIL | 27920 C>T, Orf8 I9I (VF=0.09) ** | 28854 C>T, N S194L (VF=1) | | | | |
| | Hamster# 13.1 | 3 days | 17423 A>G, Orf1b Y1319C (VF=1) | 20667 A>G, Orf1b Q2400Q (VF=0.94) | NIL | 27920 C>T, Orf8 I9I (VF=0.06) ** | 28854 C>T, N S194L (VF=1) | | | | |
| | | 5 days | 17423 A>G, Orf1b Y1319C (VF=1) | 20667 A>G, Orf1b Q2400Q (VF=0.93) | NIL | 27920 C>T, Orf8 I9I (VF=0.12)* | 28854 C>T, N S194L (VF=1) | | | | |
| | Hamster# 13.2 | 3 days | 17423 A>G, Orf1b Y1319C (VF=1) | 20667 A>G, Orf1b Q2400Q (VF=0.99) | 27147 G>T, Orf5 D209Y (VF=0.01)**[¶] | 27920 C>T, Orf8 I9I (VF=0.15) * | 28854 C>T, N S194L (VF=1) | | | | |
| | | 5 days | 17423 A>G, Orf1b Y1319C (VF=1) | 20667 A>G, Orf1b Q2400Q (VF=1) | 27147 G>T, Orf5 D209Y (VF=0.11)*[¶] | 27920 C>T, Orf8 I9I (VF=0.77) | 28854 C>T, N S194L (VF=1) | | | | |
| | Hamster# 13.3 | 3 days | 17423 A>G, Orf1b Y1319C (VF=1) | 20667 A>G, Orf1b Q2400Q (VF=0.99) | NIL | 27920 C>T, Orf8 I9I (VF=0.1) ** | 28854 C>T, N S194L (VF=0.98) | | | | |
| HK-15 | Cell culture | -- | NIL | NIL | 10981 G>T, Orf1a V3572V (VF=0.18) * | 17423 A>G, Orf1b Y1319C (VF=1) | NIL | 28854 C>T, N S194L (VF=1) | | | |
| | Hamster# 15.1 | 3 days | NIL | 7768 C>T, Orf1a I2501I (VF=0.01)**[¶] | 10981 G>T, Orf1a V3572V (VF=0.1) ** | 17423 A>G, Orf1b Y1319C (VF=1) | NIL | 28854 C>T, N S194L (VF=1) | | | |
| | | 5 days | NIL | 7768 C>T, Orf1a I2501I (VF=0.08)**[¶] | 10981 G>T, Orf1a V3572V (VF=0.14) * | 17423 A>G, Orf1b Y1319C (VF=1) | NIL | 28854 C>T, N S194L (VF=0.99) | | | |
| | Hamster# 15.2 | 3 days | 1633 A>C, Orf1a K456N (VF=0.02)**[¶] | NIL | 10981 G>T, Orf1a V3572V (VF=0.07) * | 17423 A>G, Orf1b Y1319C (VF=1) | 26125 C>A, Orf3a Q245K (VF=0.12)*[¶] | 28854 C>T, N S194L (VF=1) | | | |
| | | 5 days | 1633 A>C, Orf1a K456N (VF=0.20)*[¶] | NIL | 10981 G>T, Orf1a V3572V (VF=0.04) ** | 17423 A>G, Orf1b Y1319C (VF=1) | 26125 C>A, Orf3a Q245K (VF=0.03)**[¶] | 28854 C>T, N S194L (VF=1) | | | |
| | Hamster# 15.3 | 3 days | NIL | NIL | 10981 G>T, Orf1a V3572V (VF=0.12) * | 17423 A>G, Orf1b Y1319C (VF=1) | NIL | 28854 C>T, N S194L (VF=1) | | | |
| | | 5 days | NIL | NIL | 10981 G>T, Orf1a V3572V (VF=0.13) * | 17423 A>G, Orf1b Y1319C (VF=1) | NIL | 28854 C>T, N S194L (VF=0.99) | | | |
| HK-92 | Cell culture | -- | 1515 A>G, Orf1a H417R (VF=1) | 4093 C>T, Orf1a I1276I (VF=0.11) * | 4899 A>G, Orf1a H1545R (VF=0.21) * | 9223 C>T, Orf1a H2986H (VF=1) | 11083 G>T, Orf1a L3606F (VF=1) | 12565 G>T, Orf1a Q4100H (VF=1) | 14805 C>T, ORF1b Y446Y (VF=1) | 17247 T>C, Orf1b R1260R (VF=1) | 26144 G>T, ORF3a G251V (VF=1) |
| | Hamster# 92.1 | 3 days | 1515 A>G, Orf1a H417R (VF=1) | 4093 C>T, Orf1a I1276I (VF=0.07) ** | 4899 A>G, Orf1a H1545R (VF=0.24) * | 9223 C>T, Orf1a H2986H (VF=1) | 11083 G>T, Orf1a L3606F (VF=1) | 12565 G>T, Orf1a Q4100H (VF=1) | 14805 C>T, ORF1b Y446Y (VF=1) | 17247 T>C, Orf1b R1260R (VF=1) | 26144 G>T, ORF3a G251V (VF=1) |
| | | 5 days | 1515 A>G, Orf1a H417R (VF=1) | 4093 C>T, Orf1a I1276I (VF=0.1) * | 4899 A>G, Orf1a H1545R (VF=0.22) * | 9223 C>T, Orf1a H2986H (VF=1) | 11083 G>T, Orf1a L3606F (VF=1) | 12565 G>T, Orf1a Q4100H (VF=1) | 14805 C>T, ORF1b Y446Y (VF=1) | 17247 T>C, Orf1b R1260R (VF=1) | 26144 G>T, ORF3a G251V (VF=1) |
| | Hamster# 92.2 | 3 days | 1515 A>G, Orf1a H417R (VF=1) | 4093 C>T, Orf1a I1276I (VF=0.09) ** | 4899 A>G, Orf1a H1545R (VF=0.35) * | 9223 C>T, Orf1a H2986H (VF=0.99) | 11083 G>T, Orf1a L3606F (VF=1) | 12565 G>T, Orf1a Q4100H (VF=1) | 14805 C>T, ORF1b Y446Y (VF=1) | 17247 T>C, Orf1b R1260R (VF=1) | 26144 G>T, ORF3a G251V (VF=1) |
| | | 5 days | 1515 A>G, Orf1a H417R (VF=0.99) | 4093 C>T, Orf1a I1276I (VF=0.13) * | 4899 A>G, Orf1a H1545R (VF=0.27) * | 9223 C>T, Orf1a H2986H (VF=1) | 11083 G>T, Orf1a L3606F (VF=1) | 12565 G>T, Orf1a Q4100H (VF=1) | 14805 C>T, ORF1b Y446Y (VF=1) | 17247 T>C, Orf1b R1260R (VF=1) | 26144 G>T, ORF3a G251V (VF=1) |
| | Hamster# 92.3 | 3 days | 1515 A>G, Orf1a H417R (VF=1) | 4093 C>T, Orf1a I1276I (VF=0.11) * | 4899 A>G, Orf1a H1545R (VF=0.19) * | 9223 C>T, Orf1a H2986H (VF=1) | 11083 G>T, Orf1a L3606F (VF=1) | 12565 G>T, Orf1a Q4100H (VF=1) | 14805 C>T, ORF1b Y446Y (VF=1) | 17247 T>C, Orf1b R1260R (VF=1) | 26144 G>T, ORF3a G251V (VF=0.99) |
| | | 5 days | 1515 A>G, Orf1a H417R (VF=1) | 4093 C>T, Orf1a I1276I (VF=0.09) ** | 4899 A>G, Orf1a H1545R (VF=0.23) * | 9223 C>T, Orf1a H2986H (VF=1) | 11083 G>T, Orf1a L3606F (VF=1) | 12565 G>T, Orf1a Q4100H (VF=1) | 14805 C>T, ORF1b Y446Y (VF=1) | 17247 T>C, Orf1b R1260R (VF=1) | 26144 G>T, ORF3a G251V (VF=1) |
| HK-95 | Cell culture | -- | 241 C>T, 5' Untranslated region (5'UTR) (VF=1) | 3037 C>T, Orf1a F924F (VF=1) | 14408 C>T, Orf1b P314L (VF=1) | 23403 A>G, S D614G (VF=1) | | | | | |
| | Hamster# 95.1 | 3 days | 241 C>T, 5' Untranslated region (5'UTR) (VF=1) | 3037 C>T, Orf1a F924F (VF=1) | 14408 C>T, Orf1b P314L (VF=1) | 23403 A>G, S D614G (VF=1) | | | | | |
| | | 5 days | 241 C>T, 5' Untranslated region (5'UTR) (VF=1) | 3037 C>T, Orf1a F924F (VF=1) | 14408 C>T, Orf1b P314L (VF=1) | 23403 A>G, S D614G (VF=0.99) | | | | | |
| | Hamster# 95.2 | 3 days | 241 C>T, 5' Untranslated region (5'UTR) (VF=1) | 3037 C>T, Orf1a F924F (VF=1) | 14408 C>T, Orf1b P314L (VF=1) | 23403 A>G, S D614G (VF=1) | | | | | |
| | | 5 days | 241 C>T, 5' Untranslated region (5'UTR) (VF=1) | 3037 C>T, Orf1a F924F (VF=1) | 14408 C>T, Orf1b P314L (VF=1) | 23403 A>G, S D614G (VF=1) | | | | | |
| | Hamster# 95.3 | 3 days | 241 C>T, 5' Untranslated region (5'UTR) (VF=1) | 3037 C>T, Orf1a F924F (VF=1) | 14408 C>T, Orf1b P314L (VF=1) | 23403 A>G, S D614G (VF=1) | | | | | |

[#] Variant frequency was reported based on the Illumina sequencing results

* The frequency of genome variants was between 10% - 30%.

** The frequency of genome variants was 10% or below.

[¶] Genome variant that developed after hamster infection.

VF: Variant frequency
